# Structures of a deAMPylation complex rationalise the switch between antagonistic catalytic activities of FICD (14/96/109)

**DOI:** 10.1101/2021.04.20.440599

**Authors:** Luke A. Perera, Steffen Preissler, Nathan R Zaccai, Sylvain Prévost, Juliette M Devos, Michael Haertlein, David Ron

## Abstract

The endoplasmic reticulum (ER) Hsp70 chaperone BiP is regulated by AMPylation, a reversible inactivating post-translational modification. Both BiP AMPylation and deAMPylation are catalysed by a single ER-localised enzyme, FICD. Here we present long-sought crystallographic and solution structures of a deAMPylation Michaelis complex formed between mammalian AMPylated BiP and FICD. The latter, via its tetratricopeptide repeat domain, binds a surface that is specific to ATP-state Hsp70 chaperones, explaining the exquisite selectivity of FICD for BiP’s ATP-bound conformation both when AMPylating and deAMPylating Thr518. The eukaryotic deAMPylation mechanism thus revealed, rationalises the role of the conserved Fic domain Glu234 as a gatekeeper residue that both inhibits AMPylation and facilitates hydrolytic deAMPylation catalysed by dimeric FICD. These findings point to a monomerisation-induced increase in Glu234 flexibility as the basis of an oligomeric state-dependent switch between FICD’s antagonistic activities, despite a similar mode of engagement of its two substrates — unmodified and AMPylated BiP.

## Introduction

The endoplasmic reticulum (ER) Hsp70, BiP, dominates the chaperoning capacity of the organelle^1^. BiP’s abundance and activity are matched to the unfolded protein load of the ER at the transcriptional level, by the canonical UPR^2^, but also post-translationally^3^. BiP AMPylation, the covalent attachment of an ATP-derived AMP moiety to the Thr518 hydroxyl group, is perhaps the best-defined BiP post-translational modification. AMPylation inactivates BiP by biasing it towards a domain-docked, linker-bound ATP-like Hsp70 state and away from the domain-undocked, linker-extended ADP-like state^4–6^. As such, AMPylated BiP (BiP-AMP) exhibits high rates of substrate dissociation and is refractory to ATPase stimulation by J-domain proteins^4–6^.

BiP AMPylation inversely correlates with the ER protein folding load, increasing upon inhibition of protein synthesis^7^ and with resolution of ER stress^4^. Conversely, as ER stress mounts, inactivated BiP-AMP is recruited into the chaperone cycle by deAMPylation^4,7,8^.

A single bifunctional enzyme, FICD, is responsible for both AMPylation^4,9,10^ and deAMPylation^11–13^ of BiP. FICD is the metazoan exemplar of a family of bacterial Fic domain proteins^14^ whose canonical AMPylation activity^15–17^ is often autoinhibited by a glutamate-containing alpha helix (α_inh_)^18,19^. In FICD, the AMPylation-inhibiting Glu234 is also essential for deAMPylation^11^. Moreover, monomerisation is able to reciprocally regulate FICD’s AMPylation/deAMPylation activity, converting the dimeric deAMPylase into a monomeric enzyme with primary BiP AMPylating functionality^20^. The recent discovery that the *Enterococcus faecalis* Fic protein (EfFic) possesses deAMPylation activity which is dependent on a glutamate homologous to FICD’s Glu234^13^, suggests conservation of the catalytic mechanism amongst Fic enzymes. However, the role of Glu234 in the oligomeric state–dependent regulation of FICD’s mutually antagonistic activities remains incompletely understood.

Fic domain proteins are unrelated to the two known bacterial deAMPylating enzymes, SidD and the bifunctional GS-ATase. Both catalyse binuclear Mg^2+^-facilitated deAMPylation reactions of a hydrolytic^21^ and phosphorolytic^22^ nature, utilising a metal-dependent protein phosphatase^21^ and nucleotidyl transferase^23,24^ protein-folds, respectively. Fic proteins have a single divalent cation binding site and are evolutionarily and structurally divergent from these deAMPylases and, therefore, likely catalyse a distinct deAMPylation mechanism.

In addition to the aforementioned enzyme-based regulatory mechanism(s), there is evidence that AMPylation is also regulated by substrate availability. Cells with a constitutively monomeric FICD retain a measure of regulated BiP AMPylation^20^. FICD specifically binds and AMPylates the domain-docked ATP-state of BiP^4,20^. Client binding partitions Hsp70s away from their ATP-state, suggesting a simple mechanism for coupling BiP AMPylation to low protein folding loads. Furthermore, the finding that FICD selectively AMPylates and deAMPylates ATP-state biased BiP suggests that FICD may recognise ATP-state specific features of its substrate in a conserved binding mode, that is independent of FICD’s oligomeric-state or BiP modification status.

Here we present a structure-based approach to determine the nature of the FICD-BiP enzyme-substrate interaction, thereby elucidating the mechanism of eukaryotic deAMPylation and the basis for its regulation by an oligomerisation-based switch in FICD’s functionality.

## Results

### FICD engages AMPylated BiP and primes a Glu234-coordinated water molecule for nucleophilic attack

Mutation of the Fic motif catalytic histidine, which acts as an essential general base in the AMPylation reaction^15,16,25^, eradicates FICD’s deAMPylation activity^11^. Upon mutation of this histidine (His363Ala) FICD and BiP-AMP formed a long-lived, trapped deAMPylation complex^20^. This feature was exploited to copurify FICD and AMPylated BiP by size exclusion chromatography (SEC). A complex of otherwise wildtype dimeric FICD^H363A^ and AMPylated BiP readily crystallised, but despite extensive efforts, these crystals did not yield useful diffraction data. However, introduction of a monomerising Leu258Asp mutation and truncation of BiP’s flexible a-helical lid yielded a heterodimeric FICD^L258D-H363A^•BiP^T229A-V461F^-AMP complex (**Fig. 1a**; see methods) that crystallised and yielded two very similar sub-2 Å datasets (**Table 1**).

**Fig. 1:**
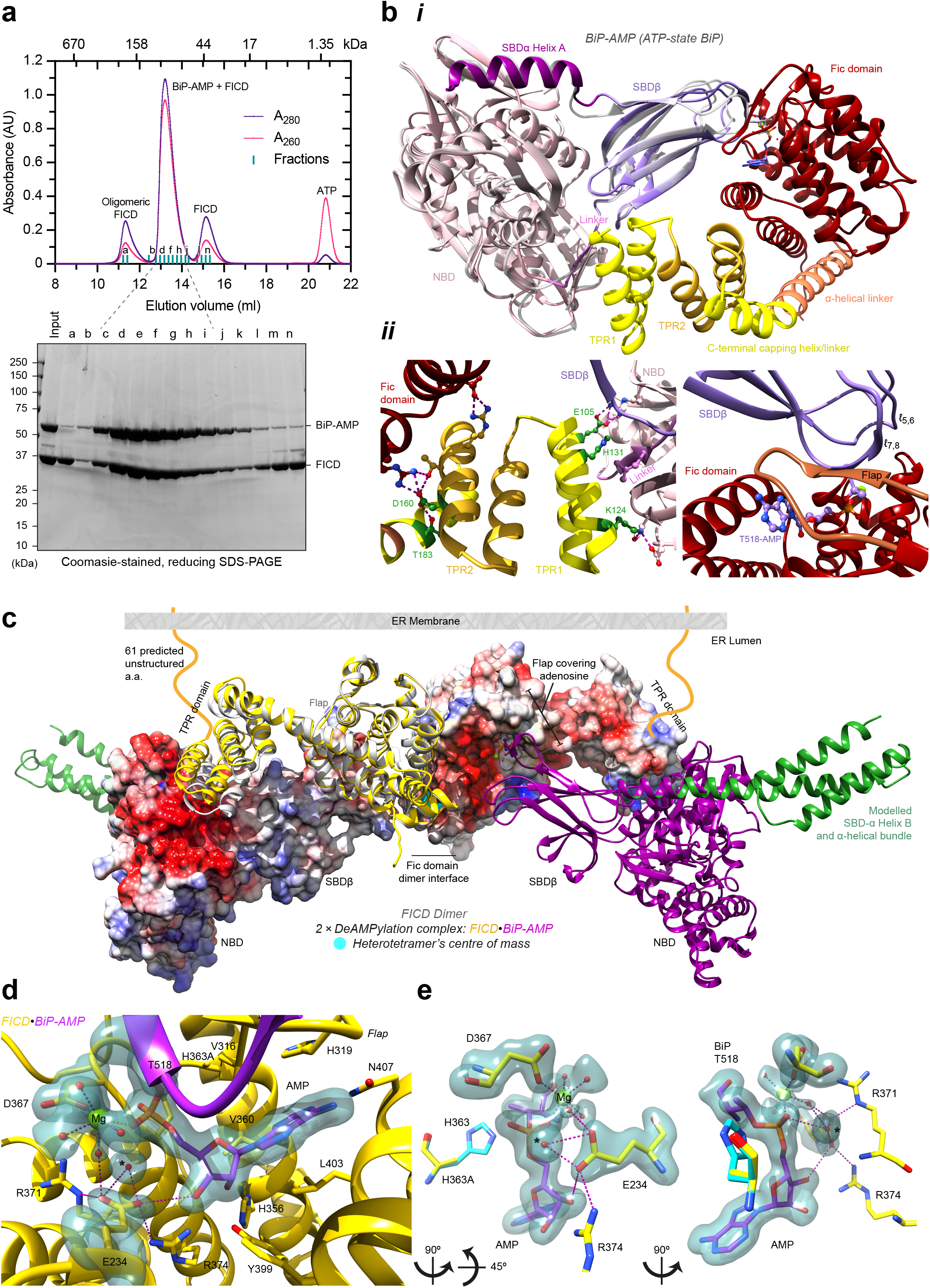
The deAMPylation complex crystal structure and mechanism of eukaryotic deAMPylation. **a,** FICD’s His363Ala mutation facilitates trapping and SEC-based copurification, of a deAMPylation complex of monomeric FICD and AMPylated BiP. **b**, The resulting deAMPylation complex crystal structure is colour-coded to illustrate its (sub)domain organisation. ***i*,** NBD-based structural superposition with the ATP-state of isolated BiP-AMP (PDB 5O4P, light grey). ***ii***, A focus on the two intermolecular interaction surfaces. Selected interdomain contacting residues are shown. Polar interactions are depicted by pink dashed lines. Residues mutated in this study are shown in green. **c**, Superposition of two heterodimeric crystal structures (purple BiPs and yellow FICDs) with an FICD dimer structure (PDB 4U0U, grey). In addition the full-length BiP lid is modelled (green) based on alignment with the BiP:ATP structure (PDB 5E84). Surfaces shown are coloured according to coulombic electrostatic potential. Note the charge complementarity between the BiP(NBD), visible on the left, and FICD(TPR1), visible on the right. For illustrative purposes the N-terminal unstructured region of FICD is shown in the context of an ER membrane. **d**, An unbiased polder-omit electron density map, contoured at 4σ, covering a region of FICD’s active site (yellow) and BiP’s Thr518-AMP (purple). Residues interacting with the AMP moiety are shown as sticks and the catalytic water is annotated with *. **e**, As in **d** but reduced to highlight Glu234’s coordination of the catalytic water molecule* in-line for nucleophilic attack into the α-phosphate. Additionally, the general acid His363 is modelled based on an alignment of PDB 6I7K.

**Table 1:** Data Collection and refinement statistics. Values in parentheses correspond to the highest-resolution shell, with the following exception: ^a^The number of molecules in the biological unit is shown in parentheses (a.u., asymmetric unit cell). ^b^The MolProbity score as a percentile is shown in parentheses, higher is better.

|  | **FICD•BiP-AMP**<br>**DeAMPylation**<br>**(State 1)** | **FICD•BiP-AMP**<br>**DeAMPylation**<br>**(State 2)** |
| --- | --- | --- |
| *Data collection* |  |  |
| **Synchrotron stations** | DLS I04-1 | DLS I04-1 |
| **Space group** | *P*2<sub>1</sub>2<sub>1</sub>2 | *P*2<sub>1</sub>2<sub>1</sub>2 |
| **Molecules in a.u.<sup>a</sup>** | 2 (2) | 2 (2) |
| **a,b,c; Å** | 95.37, 104.08,<br>105.63 | 95.00, 103.89,<br>104.79 |
| **α, β, γ; °** | 90.00, 90.00, 90.00 | 90.00, 90.00, 90.00 |
| **Resolution, Å** | 105.63-1.70<br>(1.73-1.70) | 52.40-1.87<br>(1.92-1.87) |
| **R<sub>merge</sub>** | 0.085 (1.299) | 0.087 (1.793) |
| **<I/σ(I)>** | 10.3 (1.2) | 11.9 (1.0) |
| **CC1/2** | 0.992 (0.585) | 0.999 (0.536) |
| **No. of unique reflections** | 115633 (5639) | 86247 (6270) |
| **Completeness, %** | 99.8 (99.3) | 100.0 (100.0) |
| **Redundancy** | 6.6 (6.5) | 6.6 (6.9) |
| *Refinement* |  |  |
| **R<sub>work</sub>/R<sub>free</sub>** | 0.195 / 0.221 | 0.177 / 0.231 |
| **No. of atoms (non-H)** | 7868 | 7575 |
| **Average B-factor, Å<sup>2</sup>** | 29.0 | 37.4 |
| **RMS Bond length, Å** | 0.003 | 0.003 |
| **RMS Bond angle, °** | 1.171 | 1.199 |
| **Ramachandran favoured region, %** | 98.37 | 98.49 |
| **Ramachandran outliers, %** | 0 | 0 |
| **MolProbity score<sup>b</sup>** | 0.81 (100<sup>th</sup>) | 1.04 (100<sup>th</sup>) |
| **PDB code** | 7B7Z | 7B80 |

The crystal structures displayed (identical) extensive bipartite protein-protein interfaces totalling 1366 Å^2^ (**Fig. 1b** and Supplementary **Fig. 1a**; state 1 crystal structure is shown). The deAMPylation substrate, AMPylated BiP, is in a domain-docked ATP-like state (despite having hydrolysed its bound MgATP), as reflected by the similarity with the isolated ATP-state BiP-AMP structure^5^ (**Fig. 1b(*i*)**; 1.02 Å RMSD across all 521 Cα pairs). The FICD tetratricopeptide repeat domain motif 1 (TPR1) contacted a tripartite BiP surface (695 Å^2^), comprised of its nucleotide binding domain (NBD), interdomain linker and substrate binding domain-β (SBDβ) (**Fig. 1b(*ii*)**, left panel). The second interface, by which FICD’s catalytic Fic domain engaged BiP’s SBDβ (671 Å^2^), contained an intermolecular β-sheet between BiP’s Thr518 bearing loop (*ℓ*_7,8_) and the Fic domain flap (implicated in a bacterial Fic protein AMPylation-substrate binding^16,19^). The AMP, covalently attached to BiP’s Thr518, was inserted into the Fic domain active site, with the adenosine occupying the same position as in FICD:nucleotide complexes^20,25^ (**Fig. 1b(*ii*)** right panel and **Supplementary Fig. 1b**). Contacts between the AMP moiety and the FICD active site contributed an additional 306 Å^2^ interaction surface to the deAMPylation complex.

Monomeric FICD retains deAMPylation activity^12,20^, although reduced relative to that of the dimeric enzyme^20^. Superposition of two monomeric FICD-containing deAMPylation complexes (state 1) with a dimeric FICD structure (PDB 4U0U; 2.58 Å RMSD over 334 Cα pairs across each FICD protomer), demonstrates that the heterodimeric deAMPylation crystal structure is compatible with a deAMPylation complex of dimeric FICD engaging two full-length BiP-AMP molecules (**Fig. 1c**). Furthermore, the schematised unstructured linker between the N-terminus of FICDs’ TPR domains and the ER membrane (**Fig. 1c**) illustrates that the modelled heterotetrameric structure is compatible with FICD’s presumed orientation within the ER^25,26^. Moreover, the alignment with dimeric FICD reveals intra-TPR domain movement away from FICD’s catalytic core (especially in the TPR1 motif region), which likely results from the interaction with the tripartite BiP surface.

The deAMPylation complex crystal structure contains well-resolved electron density for BiP’s AMPylated Thr518 residue within FICD’s active site (**Fig. 1d**). The phosphate of Thr518-AMP is coordinated by a Mg^2+^ held in position by FICD’s Asp367. A similarly-positioned Mg^2+^ coordinates the a and β phosphates of ATP in the AMPylation-competent enzyme^20^. Glu234 (located atop the α_inh_) tightly engages a water molecule within FICD’s oxyanion hole (**Fig. 1d** and **Supplementary Fig. 1b**). The latter Fic domain feature contributes towards the stabilisation of ATP’s a and β phosphates in the AMPylating enzyme.

The aforementioned Glu234-coordinated water molecule sits almost directly in-line with the Pα-Oγ(Thr518) phosphodiester bond (**Fig. 1e** and **Supplementary Fig. 1b**) and likely participates in catalysis. When also modelled with a catalytic histidine (from PDB 6I7K; 0.45 Å RMSD over 214 Cα pairs aligned over the Fic domain residues 213– 426) the structure is highly suggestive of an acido-basic hydrolytic mechanism of eukaryotic deAMPylation: Glu234 aligns and activates a water molecule for an S_N_2-type nucleophilic attack into the a-phosphate with His363 positioned to facilitate a concerted protonation of the Thr518 alkoxide leaving group (generating unmodified BiP and AMP as products^11^).

### The deAMPylation complex crystal structure is representative of the solution structure of dimeric FICD engaged with AMPylated BiP

To assess the validity of the structural insights gained from the heterodimeric deAMPylation complex crystal (obtained with monomeric FICD^L258D-H336A^ and a lid-truncated BiP-AMP), a solution-based structural method was employed using intact proteins. Low-resolution structures of biomacromolecules can be resolved by small angle X-ray and neutron scattering (SAXS/SANS). SAXS is sensitive to electron density, while SANS is sensitive to atomic nuclei. For mixed complexes with two components, contrast variation SANS is able to distinguish between proteins that are differentially isotopically labelled. To enable this analysis, complexes of partially deuterated and non-deuterated dimeric FICD^H363A^ and full-length BiP-AMP were copurified by SEC into buffers with varying D_2_O content. Contrast variation solution scattering data were subsequently collected (**Fig. 2a**).

**Fig. 2:**
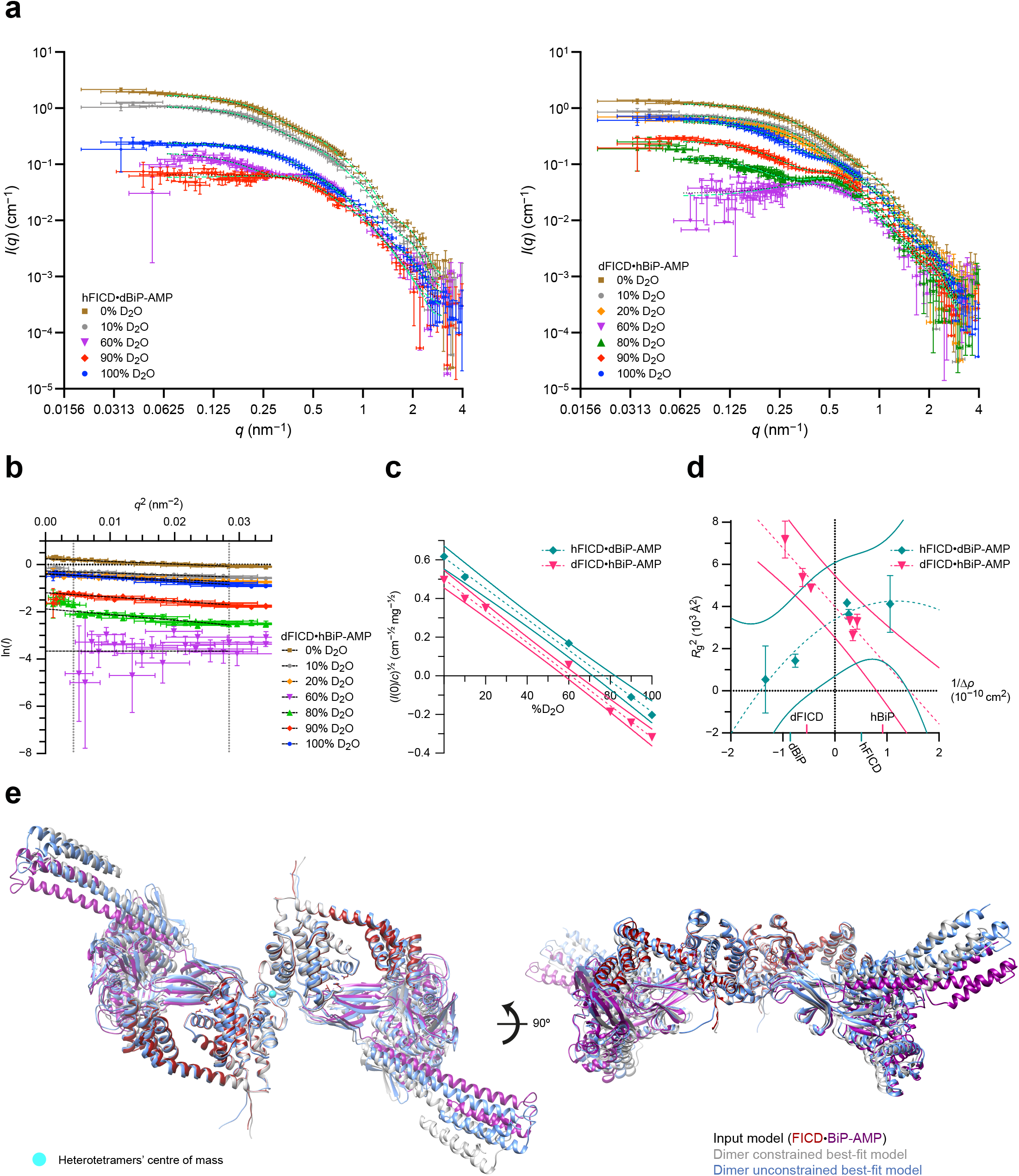
The DeAMPylation complex solution structure. **a**, Contrast-variation small angle neutron scattering (SANS) curves of copurified dimeric FICD and full-length AMPylated BiP. Overlaid dotted black lines are theoretical scattering curves based on the modelled heterotetramer shown in **Fig. 1c**, dashed green lines are the theoretical scattering curves from flex-fitting of the input heterotetramer model with a constrained FICD dimer interface. In each experiment ‘d’ and ‘h’ refer to the partially deuterated and non-deuterated component respectively. Error bars represent SEM with respect to the number of pixels used in the data averaging. **b**, Guinier plot of partially deuterated FICD and non-deuterated AMPylated BiP. **c**, Scattering amplitude plots. Linear best-fits are shown with dashed lines and 95% confidence interval bands are shown with colour-matched solid lines. **d**, Stuhrmann plot with best-fit dashed curves. 95% confidence prediction bands are shown with solid lines. The determined match points of the individual complex components are indicated on the *x*-axis. Error bars in **c** and **d** are derived from the standard errors of the Guinier fits. **e**, Optimal flex-fit structures with respect to overall agreement of theoretical scattering to all experimental contrast-variation SANS datasets. Output structures are aligned to the input heterotetramer model, itself derived by imposing the C2 symmetry of the FICD dimer (PDB 4U0U) onto the heterodimeric deAMPylation complex crystal structure as in **Fig. 1c**.

Analysis of the low-*q* Guinier region (**Fig. 2b** and **Supplementary Fig. 2a**) provided information pertaining to the forward scattering, *I*(0), and radius of gyration, *R*_g_, in each solution. The former, along with calculation of each complex’s contrast match point (CMP; **Fig. 2c**), permitted the estimation of the complex molecular weight (**Supplementary Table 1**) — which was in good agreement with a FICD•BiP-AMP 2:2 complex. The Stuhrmann plot (derived from the square of the *R*_g_ data against the reciprocal of the contrast)^27^ provided information on the internal arrangement of the heterotetramer (assigning FICD to the inside of the complex) and size (*R*_g_) of the overall complex and its constituent components; all of which are consistent with those calculated from the modelled heterotetramer structure (**Fig. 2d** and **Supplementary Table 1**).

The Stuhrmann plot’s shape provides additional information. The relatively linear Stuhrmann plot derived from the deAMPylation complex containing partially deuterated FICD, suggests that this complex has a scattering length density (SLD) centre which is very close to the complexes centre of mass (COM). The converse is true for the partially deuterated BiP complex’s Stuhrmann fit that reveals no overlap between the latter’s SLD centre and COM. As partial-deuteration of a component increases its relative contribution to the SLD, these findings are consistent with a heterotetramer in which the centre of mass lies in the plane of the FICD dimer and above the plane of the majority of the BiP mass. This arrangement fits well the structural model presented in **Fig. 1c**.

Moreover, across the scattering range and at all D_2_O concentrations, the theoretical scattering profile of the heterotetramer (modelled in **Fig. 1c**) correlated well with the observed experimental scattering, with an overall average χ^2^ of 3.4 ± 4 (mean ± SD) or 2.4 ± 2 following anomalous dataset removal (**Fig. 2a** and **Supplementary Fig. 2b**).

This was true even at D_2_O concentrations close to the CMP for each deAMPylation complex, where the scattering profile is very sensitive to both the shape and stoichiometry of the particles in solution. Furthermore, the best flex-fit structure (generated for each scattering dataset by allowing the input structure to undergo normal mode flexing of its domains) did not significantly improve model fitting. The SANS data thus indicate that the vast majority of particles in solution are engaged in a heterotetramer with neutron scattering properties predicted by a model based on the heterodimer crystal structure.

By analysing the data over the entire scattering *q*-range, through flex-fitting, it is also possible to capture some of the dynamics of the solution structure. Although no individual flex-fit structure produced a significantly reduced average χ^2^ across all datasets, a number of flex-fit output structures did have significantly different and reduced χ^2^ variance (**Supplementary Fig. 2c**, underlined). The majority of flex-fit structures possessed *R*_g_ parameters which were in good agreement with the Stuhrmann derived *R*_g_ values (**Supplementary Fig. 2d**) and the principal variation in the flex-fit structures was evident in BiP(NBD) and FICD(TPR) domain reorientation and in the BiP lid region (**Supplementary Fig. 2e–f**). Only around half of the flex-fit output structures maintained the C2 rotational symmetry present in the input heterotetramer structure (**Supplementary Fig. 2c–d**, bold), which stems from the C2 symmetry of the FICD dimer. As symmetry is expected for an average solution structure of a (symmetrical) dimeric FICD fully occupied at two independent BiP binding sites, each flex-fitting strategy yielded only best-fit structure which was both symmetrical and had a significantly reduced χ^2^ SD (**Fig. 2e**, **Supplementary Fig. 2g**, **2c–d** bold and underlined and **Supplementary Movie 1**). The best-fit structure derived from leaving the high affinity FICD dimer interface unconstrained (mean χ^2^ goodness-of-fit across the reduced data set 1.7 ± 0.4) is closer in conformation to the input structure than that obtained with a restrained dimer-interface (mean χ^2^ 2.4 ± 0.8), with an RMSD of 5.4 and 7.1 Å (across 1,892 Cα pairs), respectively. Both output structures demonstrate good *R*_g_ agreement with the Stuhrmann analysis. Importantly, the complexes’ FICD *R*_g_s are increased, and in better agreement with the experimentally derived values, relative to the input structure (**Supplementary Fig. 2d** and **Supplementary Table 1**). Therefore, the observed model deviation is indicative of additional deAMPylation complex flexibility in solution, in particular in the composite FICD(TPR)-BiP(NBD) interface and in the disposition of the BiP lid. This flexibility is inaccessible to crystallographic analysis of BiP (complexes) but is consistent with previous observations of Hsp70 conformational dynamics in the Hsp70 ATP-state^6,28^.

### Engagement of the FICD TPR domain with BiP-AMP is essential for complex assembly and deAMPylation

To test the importance of contacts between FICD’s TPR domain and BiP in complex formation, catalytically inactive (His363Ala) but structurally intact FICD variants (**Supplementary Fig. 3a–c**) were analysed for their ability to interact with immobilised BiP by BioLayer Interferometry (BLI). As FICD selectively binds to the ATP-state of BiP^20^, BiP was pre-incubated with MgATP (**Supplementary Fig. 3d**). Consistent with previous findings^20^, BiP bound more tightly to monomeric FICD^L258D-H363A^ than to dimeric FICD. The converse was true for AMPylated BiP. Complex dissociation was further accelerated by the addition of ATP to the dissociation buffer (**Fig. 3a**); via an allosteric effect on FICD, when engaging unmodified BiP:ATP, or by competition for FICD’s active site when engaging BiP-AMP^20^. However, upon removal of the TPR1 motif, dimeric FICD lost all appreciable binding to either BiP ligand. As predicted by the mode of TPR binding in the crystal structures, the isolated TPR domain measurably interacted with BiP ligands irrespective of their modification status.

**Fig. 3:**
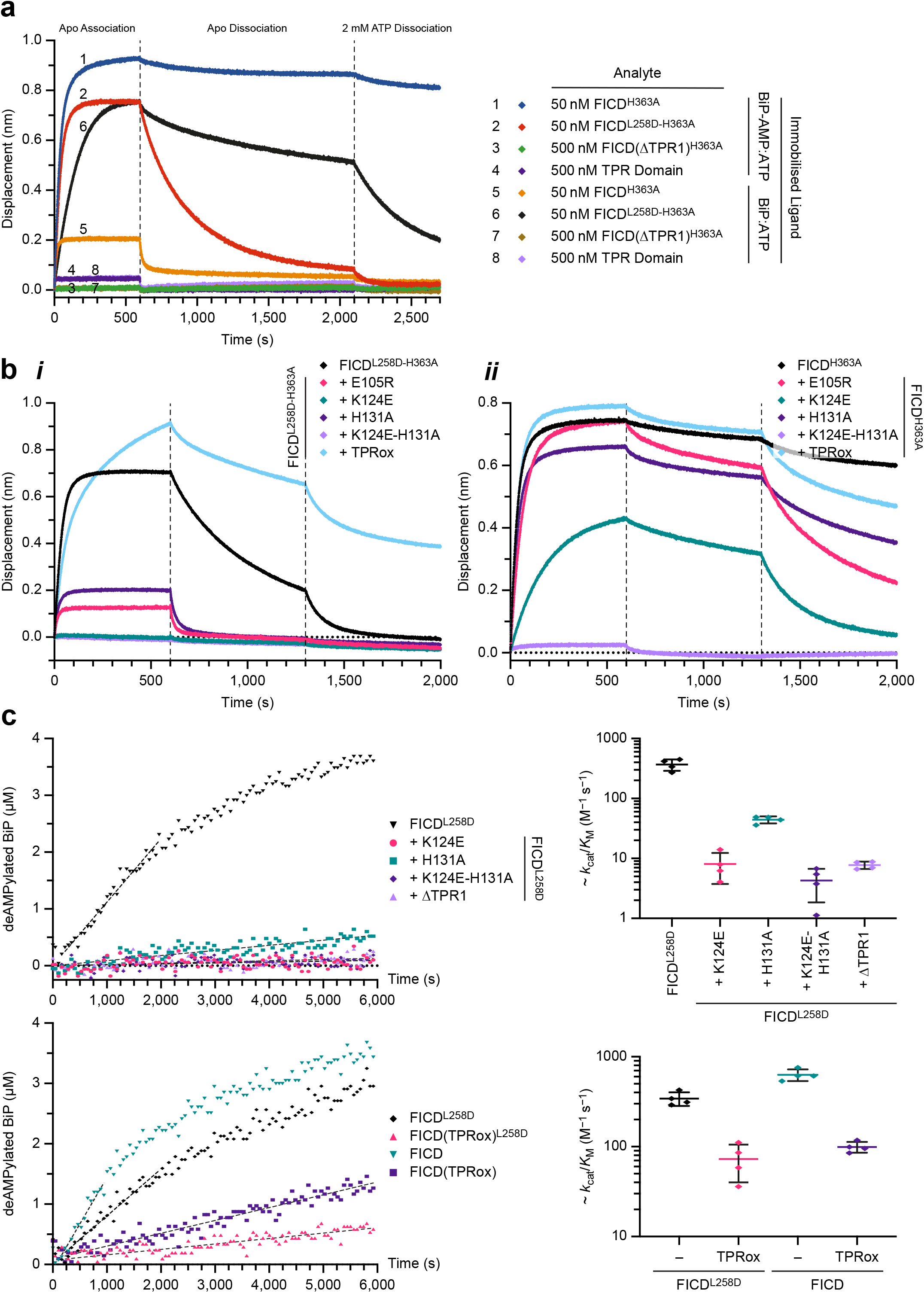
FICD’s TPR domain is essential for AMPylated BiP binding and deAMPylation. **a**, Representative BLI association-dissociation curves of FICD analytes from immobilised BiP bound to ATP (either AMPylated or unmodified), from n = 3 independent experiments. **b**, Representative BLI analysis of TPR domain mutants of monomeric (***i***) and dimeric (***ii***) FICD binding to immobilised AMPylated BiP, from n = 3 independent experiments. **c**, FP derived analysis of the ability of different FICD variants to deAMPylate BiP. Left, deAMPylation FP-derived time courses of BiP-AMP(FAM) deAMPylation. Fits of the initial linear enzyme velocities are overlaid. Right, resulting quantification of the approximate catalytic efficiencies of the different FICD variants. Mean values of approximate *k*_cat_/*K*_M_ values for each FICD variant ± SD, from n = 4 independent experiments, are shown.

The introduction of point-mutations into residues at the FICD(TPR1)-BiP interface (**Fig. 1b** and **Supplementary Fig. 1a**) significantly affected the kinetics of FICD association and dissociation of both monomeric and dimeric FICD variants (**Fig. 3b**). This agrees with the idea (supported by the solution structure) that monomeric and dimeric FICD similarly engage AMPylated BiP. Moreover, in keeping with the crystallographically observed multivalent nature of the deAMPylation complex, the kinetics of FICD^L258D-H363A^•BiP-AMP interaction appears biphasic and becomes increasingly monophasic upon disruption of FICD(TPR1)-BiP contacts (**Fig. 3b(*i*)**).

To address the role of interdomain contacts between FICD’s TPR and catalytic Fic domain in deAMPylation complex stability, one of two contacting residues within FICD’s TPR2 motif (Asp160) was mutated (**Fig. 1b(*ii*)**). However, FICD’s TPR domain has also been observed to fully disengage from the capping/linker helix, exhibiting a ‘TPR-out’ conformation (PDB 6I7K and 6I7L). To analyse the effect of perturbed interdomain contacts, whilst maintaining the BiP binding–competent ‘TPR-in’ conformation, Asp160 and Thr183 (FICD capping helix; **Fig. 1b(*ii*)**) were both mutated to cysteines and oxidised to stoichiometrically form an intramolecular disulphide bond (TPRox, **Supplementary Fig. 3c**). TPR oxidation within monomeric FICD^L258D-H363A^ resulted in more biphasic kinetics and a significant decrease in dissociation rate from BiP (**Fig. 3b(*i*)**), suggesting that the covalent fixation of the ‘TPR-in’ conformation outweighs the destabilising effects of perturbing the intramolecular Fic-TPR domain contact. Notably, the effect on dimeric FICD was less pronounced (**Fig. 3b(*ii*)**). These measurements are consistent with the fact that the ‘TPR-out’ conformation has only been observed in monomeric FICD structures^20^ and suggest that dimeric FICD has an intrinsically less flexible TPR domain. Nevertheless, TPR oxidation does alter dimeric FICD binding kinetics. The increased FICD dissociation rate, which is further exaggerated by the addition of ATP in the second dissociation phase, implicates Fic-TPR domain communication in the regulation of complex association-dissociation kinetics.

Consistent with the essential role played by the TPR domain in deAMPylation complex assembly, mutation or removal of the TPR1 motif reduced the catalytic efficiency (*k*_cat_/K_M_) of in vitro deAMPylation (**Fig. 3c**, top, and **Supplementary Fig. 3e–g**). As expected from previous analysis of FICD-mediated deAMPylation under substrate-limited conditions^20^, monomerisation of FICD was observed to diminish the rate of deAMPylation under a steady-state kinetic regime (**Fig. 3c**, bottom). Interestingly, having not significantly decreased the observed affinity for AMPylated BiP, TPR domain oxidation appreciably compromised the deAMPylation activity of both monomeric and dimeric FICD (**Fig. 3c**, bottom). This effect on catalytic efficiency presumably reflects a contribution of TPR domain flexibility or intra-FICD interdomain communication towards deAMPylation turnover number (*k*_cat_).

### FICD’s TPR domain is responsible for the recognition of unmodified ATP-state BiP

The importance of contacts between FICD’s TPR domain and BiP to deAMPylation, demonstrated above, explains previous observations that the isolated AMPylated BiP SBD is refractory to FICD-mediated deAMPylation^11^. It is noteworthy that FICD also specifically binds^20^ and AMPylates ATP-state BiP with a preference for more domain-docked BiP mutants and fails to AMPylate the isolated BiP SBD^4^. Furthermore, the observation that FICD’s interaction with unmodified BiP:ATP was abrogated by TPR1 deletion (**Fig. 3a**) hints at the possibility that FICD recognises the ATP-state of unmodified BiP (for AMPylation) in a similar fashion to ATP-state biased BiP-AMP (for deAMPylation).

Structures of unmodified BiP indicate that a domain-undocked ADP-state BiP loses the tripartite NBD-linker-SBDβ surface that is recognised by FICD’s TPR1 motif in the context of deAMPylation (**Supplementary Fig. 4a** and **Supplementary Movie 2**). Furthermore, even if FICD were able to bind the NBD or the *ℓ*_7,8_ SBDβ region (which also becomes less accessible in BiP’s ADP-state) of a nucleotide-free (apo) or ADP-bound BiP, the Hsp70’s heavy bias towards the domain-undocked conformation^6,29^ would render engagement of the other FICD-BiP interaction surface unlikely (**Supplementary Fig. 4a** and **Supplementary Movie 2**).

To test the potential role of conserved TPR-BiP contacts in formation of an AMPylation complex we returned to the BLI setup of **Fig. 3b**, but with ATP-bound unmodified BiP immobilised as a ligand. In this context the effect of TPR1 motif mutations on FICD binding were magnified relative to their effect on the deAMPylation complex (**Fig. 4a**). This is consistent with the absence of a covalently linked AMP moiety, engaging FICDs active site, increasing the relative contribution of TPR-BiP contacts to the overall complex interaction. Loss of TPR-BiP contacts by surface mutations in TPR1 also impaired BiP AMPylation by monomeric FICD in vitro (**Fig. 4b** and **Supplementary Fig. 4b**), paralleling the effect of these mutations on deAMPylation complex assembly and in vitro deAMPylation activity (**Fig. 3**). Of note, impairment of interdomain (TPR-Fic) communication by TPR oxidation, although stabilising the pre-AMPylation complex of monomeric FICD and BiP:ATP (**Fig. 4a**), decreases the in vitro AMPylation rate.

**Fig. 4:**
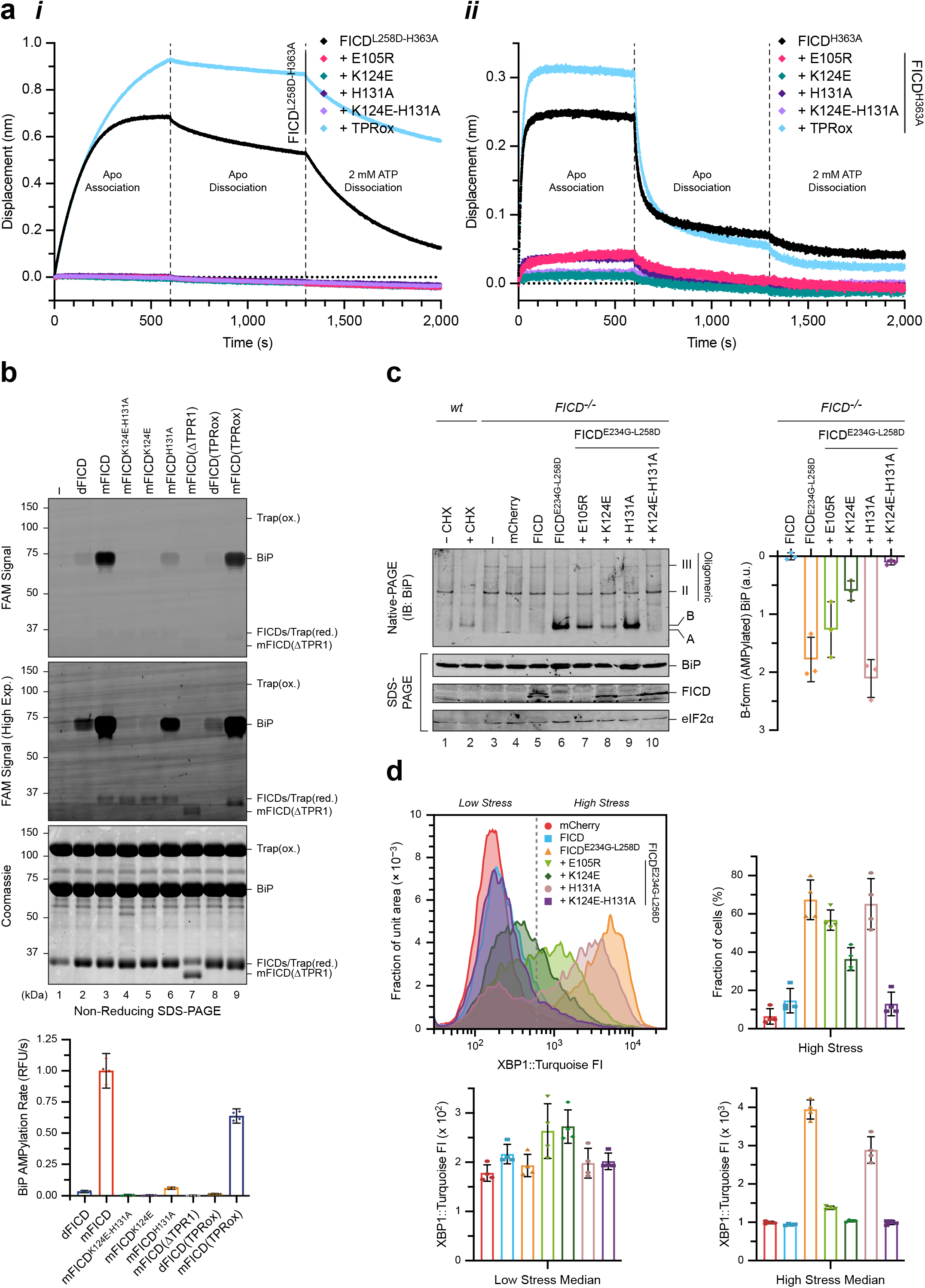
FICD’s TPR domain is essential for the recognition and AMPylation of ATP-bound BiP. **a**, Representative BLI analysis of TPR domain mutants of monomeric *(**i**)* and dimeric *(**ii**)* FICD binding to immobilised ATP-bound BiP, from n = 3 independent experiments. **b**, Fluorescence and Coomassie gel-images of an in vitro AMPylation assay, utilising ATP(FAM) as the AMPylation co-substrate, in the presence of excess product trap (Trap(ox)) to discourage BiP-AMP(FAM) deAMPylation. dFICD, dimeric FICD; mFICD, monomeric FICD^L258D^. Gels from a representative experiment are shown with the initial rates (mean ± 95% CI) of BiP-AMPylation (in relative fluorescent units/s), normalised to the rate of mFICD-mediated BiP-AMPylation, from n = 4 independent experiments. Note, the lack of correlation between FICD (*cis*)auto-AMPylation and BiP substrate AMPylation. **c**, Native-PAGE immunoblot analysis of the accumulation of AMPylated (B-form) BiP in *FICD^-/-^* CHO cells transfected with FICD variants, as indicated. Major, non-AMPylated BiP species (A, II and III) are noted. Right, quantification of AMPylated B-form BiP from n = 3 independent experiments (mean ± SD). **d**, Histograms of the FACS signal of an XBP1::Turquoise UPR reporter in *FICD^-/-^* CHO cells expressing the indicated FICD derivatives. Note the bimodal distribution of the fluorescent signal in FICD-transfected cells. Quantification of the fraction of cells that are stressed, as well as the median FACS signal of the low and high stressed cell populations are shown from n = 4 independent experiments (mean values ± SD). Bars and datapoints are (colour-)coded according to the histogram legend.

To examine the effect of the TPR surface mutations on BiP AMPylation in cells, we compared the ability of otherwise wildtype, hyperactive, monomeric FICD lacking the gatekeeper glutamate (FICD^E234G-L258D^) and TPR mutant versions thereof to promote a pool of AMPylated BiP in cells. Levels of AMPylated BiP, detected by its mobility on native-PAGE, were significantly lower in cells targeted with the FICD^K124E-E234G-L258D^ and FICD^K124E-H131A-E234G-L258D^ TPR1 mutations (**Fig. 4c**). The higher levels of expression of the TPR1 mutant FICDs (compared to FICD^E234G-L258D^) is consistent with previous observations of FICD expression levels inversely correlating with the variant’s AMPylation activity (within transiently transfected *FICD^-/-^* cells)^20^.

BiP inactivation, by deregulated AMPylation, can cause considerable ER stress^20^. This feature was exploited to quantify the functional effect of the TPR1 mutations in an orthogonal assay, based on the ER stress-responsive reporter XBP1::Turquoise, utilising flow cytometry (**Fig. 4d** and **Supplementary Fig. 4c**). In cells expressing the various TPR1 mutant FICD derivatives, reporter activity (analysed by its bimodal distribution) correlated well with the levels of AMPylated BiP detected by native-PAGE and with the hierarchy of the mutations’ effects on BiP binding (**Fig. 3b**). Together these observations lead us to conclude that TPR surface mutations in residues that contact BiP in the deAMPylation complex also contribute to enzyme-substrate interaction during FICD-induced AMPylation. Moreover, BiP’s Th518 can be readily modelled into the active site of a AMPylating monomeric FICD alongside its MgATP co-substrate, by alignment with the deAMPylation complex’s Fic domain (**Supplementary Fig. 4d**). This provides further support for there being a similar mode of FICD substrate engagement in its mutually antagonistic enzymatic activities.

### Increased Glu234 flexibility enfeebles monomeric FICD deAMPylation activity

The deAMPylation complex presented in **Fig. 1** explains the essential role of gatekeeper Glu234 in Fic domain-catalysed deAMPylation^11,13^. However, a second sub-2 Å deAMPylation complex-crystal structure, which is almost identical to that previously presented (**Table 1**, **Supplementary Fig. 5a** and **Supplementary Movie 1**), hints at an important detail. As in the state 1 structure (**Fig. 1**), the FICD active site contains clear electron densities for BiP’s Thr518-AMP, Fic domain catalytic residues and a coordinated Mg^2+^ cation (**Supplementary Fig. 5a**). However, alignment with the state 1 structure reveals a clear difference in the orientation of Glu234 (**Fig. 5a**, **Supplementary Fig. 5c** and **Supplementary Movie 3**). In the second, state 2, structure the Glu234 sidechain points further away from the position of the catalytic water molecule, that was so clearly visible in state 1, and more towards Mg^2+^.

**Fig. 5:**
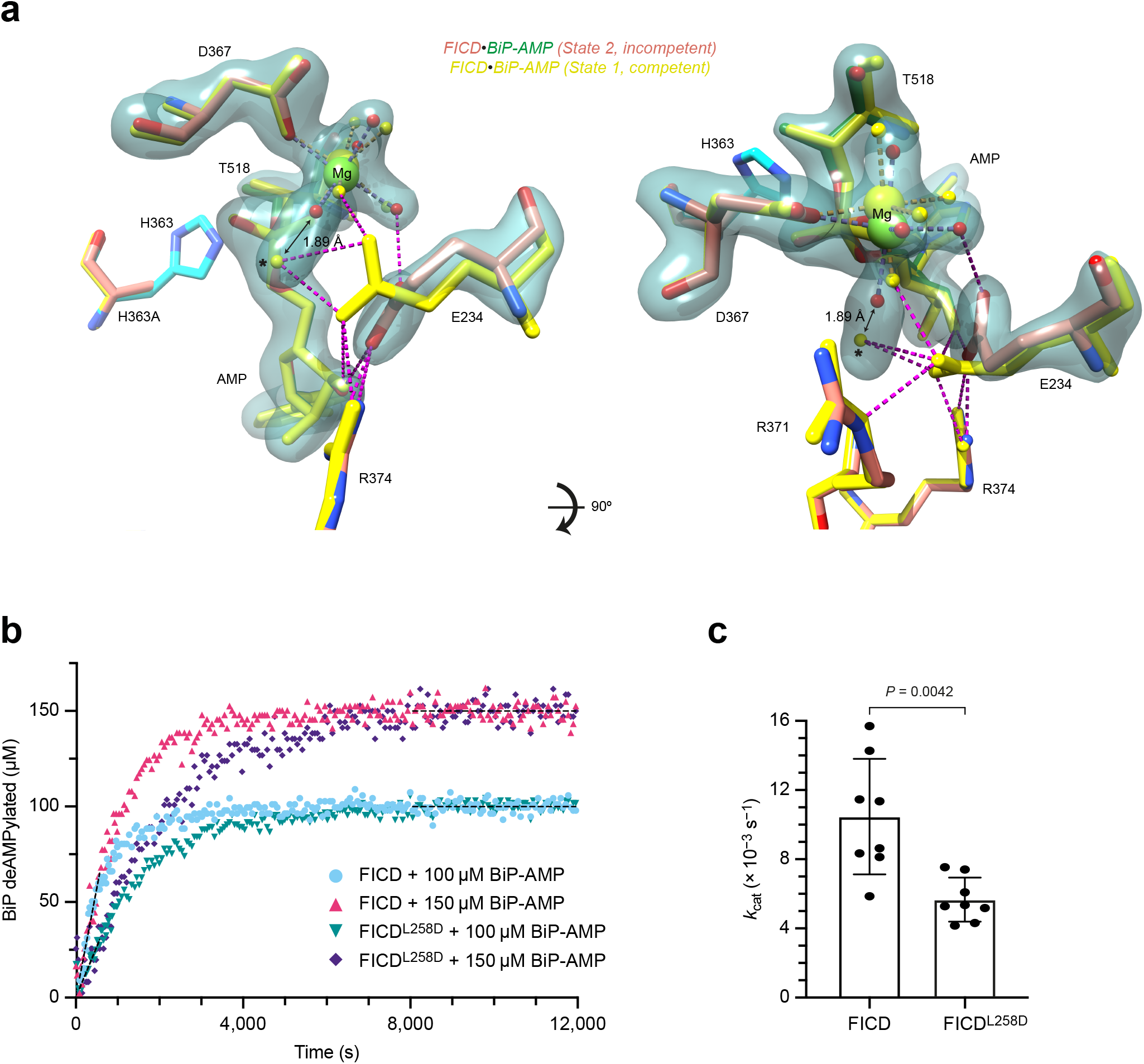
FICD monomerisation increases gatekeeper Glu234 flexibility and decreases the deAMPylation *k_cat_*. **a**, An unbiased polder-omit electron density map from a second deAMPylation complex structure (state 2), contoured at 6σ, covering Fic domain catalytic residues of particular importance (orange), the Mg^2+^-coordination complex and BiP’s Thr518-AMP (green). The reduced (state 2) active site is aligned with the active site of the (deAMPylation competent) state 1 complex (yellow). His363 is modelled from an alignment of catalytically competent FICD (PDB 6I7K, as in **Fig. 1e**). Residues interacting with the AMP moiety are shown as sticks and the catalytic water (from state 1) is annotated with *. The distance between the Mg^2+^ first-coordination sphere water (red, state 2) and the (state 1) catalytic water* is annotated. H-bonds formed by Glu234 are shown as pink-dashed lines. **b**, A representative BiP-deAMPylation time course with 10 μM FICD or FICD^L258D^, demonstrating that 100 and 150 μM BiP-AMP both represent saturating concentrations of deAMPylation substrate. **c**, The derived *k*_cat_ parameters, from n = 4 independent experiments with two saturating concentrations of BiP-AMP (as in **b**). The mean ± SD is shown with the *P*-value from a two-tailed Welch’s *t*-test annotated.

The variability in Glu234 conformation noted above fits previous observations that FICD monomerisation increases Glu234 flexibility, disfavouring autoinhibition of AMPylation activity^20^. The reorientation of Glu234 noted in state 2 also informs the deAMPylation reaction, as it results in a slight shift in the Mg^2+^ octahedral coordination complex (**Fig. 5a** and **Supplementary Movie 3**). Although there is some remaining electron density in the region of the catalytic water molecule noted in state 1, this density merged with the electron density of a Mg^2+^-coordinating water molecule. The elongated density is incompatible with the modelling of two water molecules (accommodating the Mg^2+^-coordination geometry requirements would necessitate an infeasible inter-water distance of 1.89 Å) and suggests that there may be a dynamic shuttling of a water to and from the primary Mg^2+^-coordination sphere into a position more conducive to catalysis. It is clear that the Glu234 position observed in the state 2 crystal structure does not permit the stable positioning of a catalytic water molecule inline for nucleophilic attack.

A corollary of the two tenets, that Glu234 is necessary for coordinating a catalytic water molecule for deAMPylation and that Glu234 flexibility increases upon monomerisation, is the prediction that FICD deAMPylation activity should decrease upon monomerisation. This has already been demonstrated in terms of a 46% decrease in catalytic efficiency (**Fig. 3c**) — the calculated *k*_cat_/*K*_M_ of FICD (630 ± 50 s^−1^ M^−1^, mean ± SEM) is 1.9-fold greater than that of FICD^L258D^ (340 ± 30 s^−1^ M^−1^). Moreover, dimeric FICD’s *k*_cat_/*K*_M_ is in good agreement with that derived from a previous Michaelis-Menten analysis of a GST-tagged FICD (600 ± 100 s^−1^ M^−1^, best-fit ± SE)^11^.

However, an increase in Glu234 flexibility is expected to intrinsically affect deAMPylation catalysis and lower the *k*_cat_. In order to directly measure the turnover number for monomeric and dimeric FICD both enzymes must be saturated with deAMPylation substrate. It was found that the initial rates of deAMPylation were indistinguishable at initial substrate concentrations of 100 and 150 μM BiP-AMP (**Fig. 5b** and **Supplementary Fig. 5d–e**), implying that FICD and FICD^L258D^ are saturated by BiP-AMP. Therefore, at these substrate concentrations the initial deAMPylation rates represent maximal enzyme velocities, from which a *k*_cat_ parameter can be extracted (**Fig. 5c**). As expected for the less-flexible Glu234-bearing dimeric FICD, its deAMPylation *k*_cat_ ({10 ± 1} × 10^−3^ s^−1^, mean ± SEM) was significantly greater (1.8-fold) than that of monomeric FICD^L258D^ ({5.7 ± 0.4} × 10^−3^ s^−1^).

Together, the comparison of deAMPylation catalytic efficiencies and turnover numbers between dimeric and monomeric FICD, suggests that the major effect of monomerisation on the kinetics of deAMPylation is mediated through a decrease in *k*_cat_. Thus, despite the apparent differences between monomeric and dimeric FICD in their affinity for BiP-AMP (**Fig. 3a**), any differences in *K*_D_ must be compensated for by the variation in *k_cat_* values — resulting in very similar *K*_M_ values of 16 ± 2 μM for dimeric and 17 ± 2 μM for monomeric FICD (mean ± SEM). Note, the *k*_cat_ and *K*_M_ values derived for dimeric FICD are in good agreement with those previously obtained from Michaelis-Menten analysis of GST-FICD: *k*_cat_ {9.9 ± 0.9} × 10^−3^ s^−1^ and *K*_M_ 16 ± 3 μM (best-fit ± SE)^11^, adding credibility to the *k*_cat_/*K*_M_ and *K*_M_ determinations.

## Discussion

Here, we have leveraged insights from crystal structures of a deAMPylation complex (the first such structure to our knowledge) of FICD and BiP-AMP to gain a detailed understanding of eukaryotic deAMPylation and a broad understanding of the enzyme-substrate interactions of FICD that underpin its mutually antagonistic activities of BiP AMPylation and deAMPylation. Biochemical and cellular studies of structure-guided mutations in FICD have shed light on both substrate level and enzyme-level regulation of BiP’s AMPylation cycle as it matches BiP activity to ER stress in a post-translational strand of the UPR (**Fig. 6**).

**Fig. 6:**
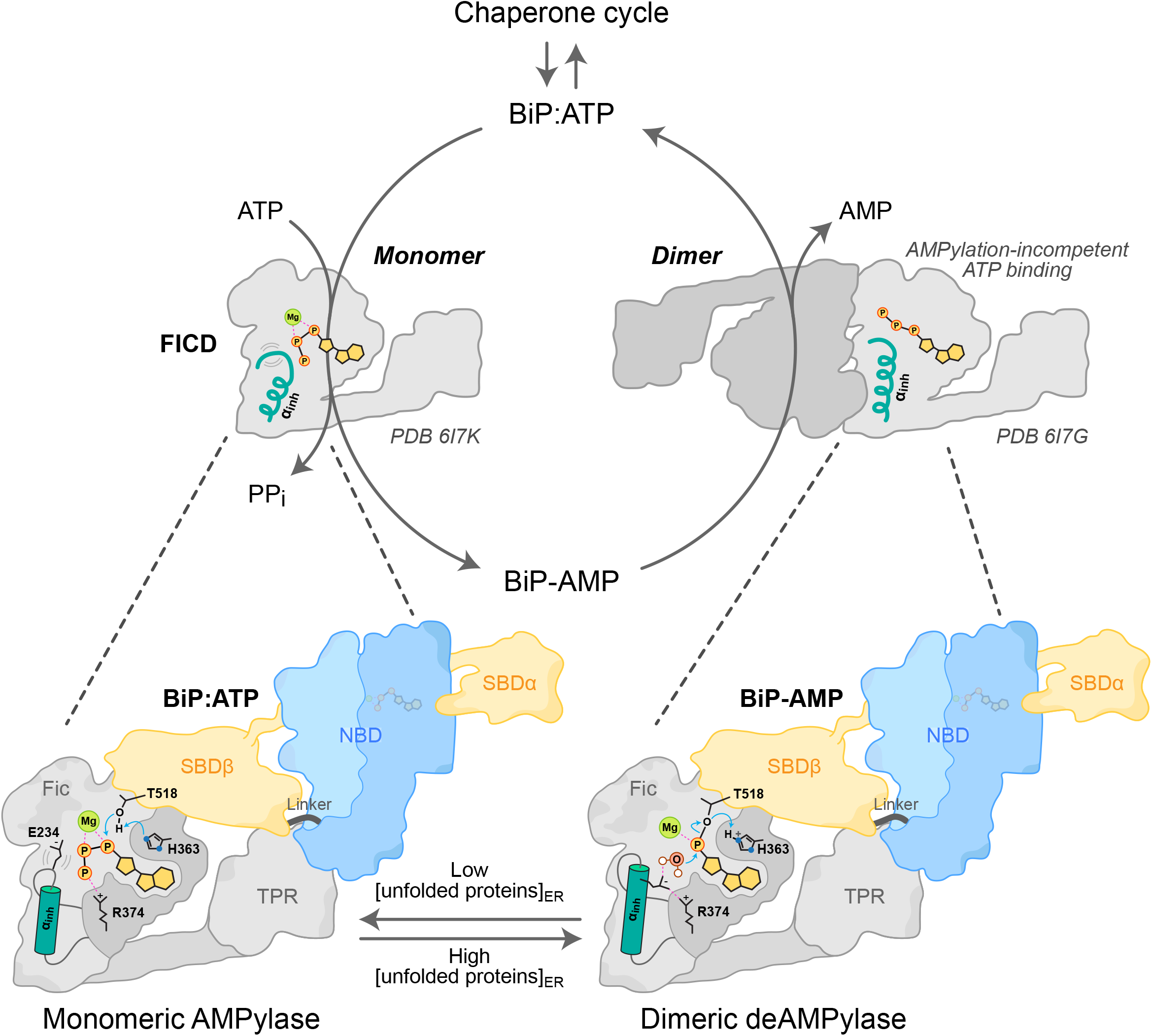
Model of FICD AMPylation and deAMPylation of BiP. FICD recognises (AMPylated or unmodified) BiP’s linker-docked NBD and the *ℓ*_7,8_ region of the SBDβ, via its TPR and catalytic domain, respectively. This is only possible when BiP is in a domain-docked ATP-like state. Dimeric FICD has a relatively rigid gatekeeper Glu234 which facilitates efficient alignment of an attacking water for BiP deAMPylation whilst prohibiting AMPylation competent binding of ATP. Conversely, monomeric FICD has a more flexible Glu234 which decreases its deAMPylation efficiency whilst permitting AMPylation competent binding of MgATP. The FICD monomer-dimer equilibrium is adjusted in response to changing levels of unfolded proteins within the ER by a yet-to-be discovered process.

The specific recognition of ATP-state BiP is mediated by an interaction of FICD’s TPR1 domain with a tripartite ATP state–specific Hsp70 surface composed of BiP’s NBD, linker and SBDβ. Moreover, the TPR domain of FICD is only able to direct BiP’s *ℓ*_7,8_ SBDβ region into the Fic domain active site when BiP’s NBD and SBD are closely opposed, as in the domain-docked ATP-state. These features explain the finding that the client protein-bound ADP-state BiP is not a substrate for AMPylation^4^ and suggests a facile mechanism for substrate-level regulation of BiP AMPylation — in which substate availability is inversely proportional to the unfolded protein load in the ER.

A reciprocal mechanism for substrate-level regulation of deAMPylation is unlikely, as AMPylated BiP is intrinsically biased towards the ATP-like domain-docked state^5^. Thus, evidence from biochemical and cell-based experiments for similar engagement of BiP in FICD-mediated AMPylation and deAMPylation, suggests that regulatory changes in FICD’s active site contribute to the enzyme’s ability to respond to changes in the burden of ER unfolded proteins. Previous studies uncovered a role for a monomerisation-induced increase in Glu234 flexibility, which permits AMPylation competent binding of MgATP within the FICD active site^20^. However, the basis for the relationship between oligomeric state and deAMPylation activity remained obscure, awaiting clarification of the enzymatic mechanism and the essential role played by Glu234 in FICD-mediated deAMPylation.

The crystal structures presented in this work provide strong support for a mechanism of eukaryotic deAMPylation that is acido-basic in nature and in which Glu234 aligns a catalytic water molecule in-line for nucleophilic attack into a-phosphate of Thr518-AMP (**Supplementary Fig. 6**). Glu234, may act as a catalytic base but through a mechanism involving late proton transfer analogous to the role played by the catalytic aspartates of some protein kinases^30,31^. This proposed deAMPylation mechanism (which also rationalises the essential role for a divalent cations and His363) is far removed from the binuclear metal-catalysed reactions catalysed by the other two known (bacterial) deAMPylases^21,24^. Moreover, other mechanisms of phosphodiester bond cleavage, including anchimeric assistance or an E1cB-type elimination reaction, which are capable of generating the products of FICD-mediated deAMPylation (AMP and unmodified BiP), are rendered extremely unlikely by the structure of the deAMPylation complex (**Supplementary Fig. 1b**).

A hydrolytic SN2-type acido-basic catalysed nucleophilic-substitution reaction, facilitated by Glu234 and His363, represents a highly plausible deAMPylation mechanism that is supported by the structure of the deAMPylation complex. As a bacterial Fic protein (EfFic) has also been observed to possess gatekeeper glutamate-dependent deAMPylation activity^13^, it is likely that the mechanism of deAMPylation outlined above is conserved across this class of proteins. This conclusion, pertaining to the immediate role of Glu234 in enabling BiP-AMP hydrolysis, permits various inferences to be made about the role of monomerisation and increased Glu234 flexibility^20^ in the regulation of deAMPylation activity. These, are supported by the direct observation of a monomeric FICD-deAMPylation complex with an alternative Glu234 conformation, resulting in a (state 2) deAMPylation non-competent active site lacking a stably coordinated catalytic water molecule. Thus, increased Glu234 flexibility, induced by FICD monomerisation, not only considerably increases AMPylation activity but also decreases the deAMPylation *k*_cat_ (**Fig. 6**).

Oligomeric-state changes in the disposition of the gatekeeper Glu234 may not be the only mechanism for enzyme-based regulation of the BiP AMPylation-deAMPylation cycle. Observations that monomeric FICD binds more tightly to unmodified BiP than BiP-AMP and the converse being true for dimeric FICD, remain unexplained by the structure of the FICD deAMPylation complex, suggesting that other factors may contribute to regulation. For example, there may well be subtle differences in the interactions between FICD and BiP mediated by changes in oligomeric state/modification status or by FICD protein dynamics; as hinted at by the crystallographic and SANS-based evidence for TPR domain flexibility and by the effects of TPR fixation on enzyme-substrate complex formation and catalysis.

These caveats notwithstanding, this study advances our mechanistic understanding of the reciprocal-regulation of enzymatic activity afforded by FICD’s oligomerisation-state dependent switch (**Fig. 6**). This leaves unanswered the question of how the FICD monomer-dimer equilibrium responds to changing conditions in the ER. There is some evidence that FICD may respond to the energy-status of the ER, as a proxy for ER stress^20^. Given that Hsp70 proteins can directly modulate the oligomeric status (and thus activity) of their own regulators within the ER^32^ and cytosol/nucleus^33^, the possibility of an additional layer of BiP-driven FICD-regulation is therefore an intriguing one to consider.

## Materials and Methods

### Plasmid construction

The plasmids used in this study have been described previously or were generated by standard molecular cloning procedures and are listed in **Supplementary Table 2**.

### Protein purification

All proteins were purified using the method for FICD protein expression detailed in^20^, with only minor modifications. In brief, proteins were expressed as N-terminal His_6_-Smt3 fusion constructs from either pET28-b vectors (expressed in T7 Express *lysY/I^q^* (NEB) *Escherichia coli* (*E. coli*) cells), or pQE30 vectors (expressed in M15 *E. coli* cells (Qiagen)). T7 Express cells were grown in LB medium containing 50 μg/ml kanamycin. M15 cells were grown in the same medium supplemented with an additional 100 μg/ml ampicillin. All cells were grown at 37 °C to an optical density (OD_600nm_) of 0.6 and then shifted to 18 °C for 20 min, followed by induction of protein expression with 0.5 mM isopropylthio β-D-1-galactopyranoside (IPTG). Cells were harvested by centrifugation after a further 16 h at 18 °C.

Only the predicted structured regions of human FICD were expressed (residues 104–445). For ‘full-length’ BiP constructs, that is to say constructs containing the complete structured region of the SBDa lid subdomain, residues 27–635 of Chinese hamster BiP were expressed. This excludes an unstructured acidic N-terminal region and the C-terminal unstructured region bearing the KDEL. Note, in the recombinantly expressed residue range hamster and human BiP are identical in terms of amino acid identity. For use as an immobilised BLI ligand full-length BiP was expressed with an avi-tag inserted C-terminal to Smt3 and N-terminal to a GS linker and hamster BiP residues 27–635.

All BiP constructs used in this study were made ATPase^34^ and substrate-binding^35^ deficient via introduction of Thr229Ala and Val461Phe mutations, respectively. Thr229Ala allows BiP to bind and domain-dock in response to MgATP, even when immobilised via an N-terminal biotinylated Avi-tag^20^. The lack of ATP hydrolysis enables BiP to remain bound to ATP in its domain-docked state for prolonged periods of time, a feature which favours binding to^20^ and AMPylation by FICD^4^. Both Thr229Ala (in the presence of ATP) and Val461Phe (independent of nucleotide) disfavour the binding of proteins within BiP’s SBD (which principally occurs in the apo or ADP-state).

Following harvesting and lysis of the bacterial pellets, proteins were purified through the use of Ni-NTA agarose (Thermo Fisher), on-bead Ulp1 cleavage, anion exchange and gel filtration chromatography as described in^20^ with minor modifications. All purification was conducted at 4 °C. Unless otherwise specified (below) anion exchanges were conducted using a RESOURCE Q 6 ml column (GE Healthcare) with a linear gradient ranging from 95% AEX-A (25 mM Tris-HCl pH 8.0) and 5% AEX-B (25 mM Tris-HCl, 1 M NaCl) to 50% AEX-A and 50% AEX-B (see **Supplementary Table 2**). Gel filtration was conducted, depending on protein size and amount, on either a HiLoad 16/60 Superdex 75 or 200 prep grade column or a S200 or S75 Increase 10/300 GL column (see **Supplementary Table 2**). All proteins were purified to homogeneity and > 95% purity, as assessed by Coomassie-stained SDS-PAGE. Unless the protein was deliberately oxidised they were supplemented after gel filtration with 1 mM tris(2-carboxyethyl)phosphine (TCEP). Proteins were concentrated to > 150 μM using centrifugal filters (Amicon Ultra; Merck Millipore), aliquoted and snap-frozen and stored at −80 °C. All protein concentrations were calculated using A_280_, measured on a NanoDrop One (Thermo Fisher), and the protein’s predicted extinction coefficient at 280 nm (ε_280_).

#### Preparative BiP AMPylation

In the case of preparative scale AMPylation of BiP, this was achieved post-Ulp1 cleavage by addition of 10 mM MgCl_2_, 5 mM ATP and 1/50 (*w/w*) GST-TEV-FICD^E234G^ (UK1479; purified as previously^20^). The AMPylation reaction was incubated for 16 h at 25 °C. GST-TEV-FICD was then depleted by a 1 h incubation with GSH-Sepharose 4B matrix (GE Healthcare). AMPylation was confirmed as being stoichiometric by intact-protein mass spectrometry (LC-ESI-MS) as previously detailed^5^.

#### Disulphide-linked FICD dimers and BiP biotinylation

Disulphide-linked FICD dimers (_s-s_FICD^A252C-H363A-C421S^; UK2269), used as a BiP-AMP trap for in vitro AMPylation assays, were oxidised and purified as in a previous study^20^. Likewise, in vitro biotinylation of N-terminally avi-tagged BiP was conducted and the proteins made apo and purified as previously described^20^ with the exception that ion-exchange fractions were diluted with glycerol and stored at −20 °C in a final buffer of TNTG (12.5 mM Tris-HCl pH 8.0, ~ 150 mM NaCl, 0.5 mM TCEP and 50% *(v/v)* glycerol), without additional gel filtration (see **Supplementary Table 2**).

#### FICD TPR domain oxidation

Purification of TPR domain oxidised (TPRox) FICD^D160C-T183C-C421S^-derivative proteins was achieved as above (for other FICDs), with the addition of an oxidation and clean-up AEX step. Note, the cysteine free FICD^C421S^ mutation was previously observed to have no effect on FICD-mediated deAMPylation or BiP-AMP binding and a slight stimulatory effect on FICD-mediated AMPylation^20^.

In order to form the disulphide-bond, the FICD protein (post-Ulp1 cleavage and Ni-NTA column elution) was diluted down to a concentration of 5 μM in a final buffer of 25 mM Tris-HCl pH 8.0 and 100 mM NaCl, supplemented with 0.5 mM CuSO_4_ and 1.75 mM 1,10-phenanthroline (Sigma), and incubated for 16 h at 4 °C. The oxidation reaction was then quenched by the addition of 2 mM EDTA. The protein solution, diluted with 25 mM Tris pH 8.0 to a final NaCl concentration of 50 mM, was then purified on a HiTrap 5 ml Capto Q column (equilibrated in 95% AEX-A and 5% AEX-B buffer) using a linear gradient of 5–50% AEX-B over 10 column volumes. Proteinaceous fractions were further purified as detailed above (beginning with RESOURCE Q column purification), culminating in the purification of dimeric or monomeric FICD (as appropriate) by gel filtration.

Stoichiometric disulphide bond formation was confirmed by the use of an electrophoretic mobility assay (see **Supplementary Fig. 3c**), in which the putatively oxidised protein was heated for 10 min at 70 °C in SDS-Laemmli buffer ± DTT; all available thiols were then reacted with a large excess of PEG 2000 maleimide (30 min at 25 °C). All unreacted maleimides were then quenched by the addition of a molar excess of DTT before samples were analysed by SDS-PAGE. Significant PEG modification of FICD(TPRox) proteins was only observed in samples first denatured in reducing conditions (+ DTT), suggesting that the two TPR domain-cysteines were not accessible for alkylation in the absence of DTT (on account of being oxidised to form an intramolecular disulphide bond).

### Protein crystallisation and structure determination

Monomeric FICD^L258D-H363A^ (residues 104–445) [UK2093] and monomeric lid-truncated BiP^T229A-V461F^-AMP (residues 27–549) [UK2090] were purified as above and gel filtered into a final buffer of T(10)NT (10 mM Tris-HCl pH 8.0, 150 mM NaCl and 1 mM TCEP). As outlined in the text, FICD’s His363Ala mutation facilitates a stable trapping of its deAMPylation substrate. As mentioned above, BiP^T229A-V461F^ favours its monomeric ATP-state, in which it is less likely to bind substrates in its SBD and to form BiP oligomers. The removal of all but helix A of the SBDa (BiP residues 27–549) was also implemented to reduce the affinity of BiP substrate binding and oligomerisation and to increase the likelihood of crystallisation and high resolution diffraction by removal of the flexible SBDα helix B, which in other Hsp70s has been documented to only transiently interact with the NBD in the ATP-state^28^. Heterodimer copurification was achieved by mixing FICD^L258D-H363A^ and BiP^T229A-V461F^-AMP in a 1.5:1 molar ratio, supplemented with an additional 250 μM ATP, 50 mM KCl and 2 mM MgCl_2_. The mixture was incubated for 10 min at 4 °C and purified by gel filtration on an S200 Increase 10/300 GL column equilibrated in TNKMT buffer (10 mM Tris-HCl pH 8.0, 100 mM NaCl, 50 mM KCl, 2 mM MgCl_2_ and 1 mM TCEP) with ≤ 5 mg of protein injected per SEC run. Heterodimeric protein fractions were pooled (as indicated in **Fig. 1a**) and concentrated to 10.3 mg/ml using a 50 kDa MWCO centrifugal filters.

Crystallisation solutions, consisting of 100 nl protein solution and 100 nl crystallisation reservoir solution, were dispensed using a mosquito crystal (SPT Labtech) and the complex was crystallised via sitting drop vapour diffusion at 25 °C. State 1 crystals were obtained from reservoir conditions of 0.1 M MES pH 6.5, 10% PEG 4000 and 0.2 M NaCl; state 2 crystals were obtained from conditions of 0.1 M Tris pH 8.0 and 25% PEG 400. Crystals were cryoprotected in a solution consisting of 25% glycerol and 75% of the respective reservoir solution (*v/v*).

Diffraction data were collected from the Diamond Light Source at 100 K (beamline I04-1), and the data processed using DIALS^36^ (state 1 crystal) or xia2^37^ (state 2 crystal) and the CCP4 module Aimless^38,39^. Structures were solved by molecular replacement using the CCP4 module Phaser^38,40^. AMPylated BiP (PDB 5O4P) and monomeric FICD (PDB 6I7L) structures from the Protein Data Bank were used as initial search models. Manual model building was carried out in COOT^41^ and refined using refmac5^42^ with TLS added. Metal binding sites were validated using the CheckMyMetal server^43^. Polder (OMIT) maps were generated using the Polder Map module of Phenix^44,45^. Structural figures were prepared using UCSF Chimera^46^, estimates of interaction surface areas were derived from PISA analysis^47^, interaction maps (**Supplementary Fig. 1**) were based on an initial output from LigPlot+^48^ and the chemical reaction pathway (**Supplementary Fig. 6**) was created in ChemDraw (PerkinElmer Informatics).

### Contrast Variation Small Angle Neutron Scattering

Non-deuterated BiP^T229A-V461F^-AMP (residues 27-635) and FICD^H363A^ (residues 104-445) [hBiP-AMP and hFICD] were purified as detailed above but were gel filtered into a final buffer of TNKMT(0.2) [TNKMT buffer with TCEP reduced to 0.2 mM]. The matchout deuterium labelled protein equivalents were produced in the ILL’s deuteration laboratory (Grenoble, France). Proteins were expressed from *E. coli* BL21 Star (DE3) cells (Invitrogen) that were adapted to 85% deuterated Enfors minimal media containing unlabelled glycerol as carbon source, as described previously^49,50^, in the presence of kanamycin at a final concentration of 35 μg/ml. The temperatures at which the cells produced the highest amount of soluble matchout-deuterated BiP or FICD were chosen for cell growth using a high cell density fermentation process in a bioreactor (Labfors, Infors HT). For BiP expression, cells were grown using a fed-batch fermentation strategy at 30 °C to an OD_600_ of 20. The temperature was then decreased to 18 °C and protein expression was induced by addition of 1 mM IPTG. After a further 22 h of protein expression at 18 °C, bacteria were harvested by centrifugation. FICD expression was conducted likewise, but with induction at OD_600_ 19 and at a temperature of 22 °C. FICD expressing cells were incubated for a further 21.5 h at 22 °C before harvesting. Matchout-deuterated proteins (dBiP^T229A-V461F^-AMP and dFICD^H363A^) were isolated and purified from deuterated cell pastes using H_2_O-based buffer systems, as mentioned above, and gel filtered into TNKMT(0.2).

Heterotetrameric complexes were copurified by gel filtration of a mixture of either dBiP-AMP and hFICD or hBiP-AMP and dFICD (in a 1.25:1 molar ratio of BiP-AMP:FICD), with ≤ 5 mg of protein injected per SEC run, supplemented with 250 μM ATP. The gel filtration was conducted on an S200 Increase 10/300 GL column equilibrated with TNKMT(0.2) buffer. Heterotetrameric complex fractions were collected and concentrated to > 7 mg/ml. Some of this purified complex was further exchanged by the same SEC process into TNKMT(0.2) in which the solvent used was D_2_O. That is to say, the complex was exchanged into 100% D_2_O buffer. Protein fractions in 100% D_2_O buffer were subsequently concentrated to > 6 mg/ml. The elution profile appeared largely identical in both deuterated and non-deuterated buffers. Complexes at different %D_2_O were obtained by either dilution with the appropriate matched buffer (± D_2_O) or by the mixing of one complex purified in 0% D_2_O buffer with the same complex in 100% D_2_O buffer.

SANS data were collected from a total of 17 samples at various D_2_O buffer compositions at 12 °C at the ILL beamline D11. Protein complexes (ranging from 4.3 to 5.5 mg/ml) were analysed in a 2 mm path-length quartz cell with a 5.5 Å wavelength neutron beam at distances of 1.4, 8 and 20.5 m. Data from relevant buffer-only controls were also collected with similar data collection times and subtracted from the radially averaged sample scattering intensities to produce the *I*(*q*) against *q* scattering curves presented in **Fig. 2a**. Scattering data were initially processed with the GRASP (Graphical Reduction and Analysis SANS Program for Matlab; developed by Charles Dewhurst, ILL) and with the Igor Pro software (WaveMetrics) using SANS macros^51^. Data analysis was conducted using Prism 8.4 (GraphPad) and PEPSI-SANS (for fitting of theoretical scattering curves and flex-fit model generation; software based on PEPSI-SAXS^52^).

Comparison of the *ln*(*Transmission*) of the 0% and 100% D_2_O buffers alone with the *ln*(*Transmission*) of each sample (not shown) confirmed that the %D_2_O of each sample was within the margin of error of the theoretical D_2_O content^53^.

Parameters from the Guinier plots were derived from fitting of the Guinier approximation^54^:

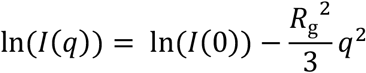

The upper and lower *q* limits for fitting are shown (grey vertical dashed lines in **Fig. 2b** and **Supplementary Fig. 2a**, except for the fitting of hFICD•dBiP-AMP in 60% D_2_O buffer where the lower *q* limit is denoted by purple vertical dashed line) and result in *qR*_g_ < 1.3 (with the exception of the fitting of dFICD•hBiP-AMP in 80% D_2_O buffer data where *qR*_g_ = 1.4).

The contrast match point analysis (CMP) in **Fig. 2c** indicated complex match points of 76.7% D_2_O (95% CI: 71.5 to 82.4% D_2_O) and 61.4 D_2_O (95% CI: 57.4 to 65.5% D_2_O) for hFICD•dBiP-AMP and dFICD•hBiP-AMP, respectively. Comparison of the experimental CMPs with theoretical values calculated by MULCh^55^ (which takes into account buffer composition effects (at 20 °C) and protein sequence, whilst assuming a 1:1 complex and 95% labile H/D-exchange) suggested that there was 66.5% deuteration of dBiP-AMP and a 63.8% deuteration of dFICD. Note, these deuteration values are less than the theoretical maximum which could have been obtained from the 85% deuterated E. coli growth media, see above. These values of (non-labile) protein (partial) deuteration were used to calculate theoretical *I*(0)/*c* values in SASSIE^56^, using the same assumptions as above. Comparison of the theoretical *I*(0)/*c* values with those determined from the experimental Guinier analysis facilitated experimental protein-complex MW estimation^57^ (**Supplementary Table 1**). The contrast at each %D_2_O (the difference in scattering length density (SLD), Δ*ρ*, between the *ρ*_protein_ and *ρ*_buffer_) was also derived from MULCh.

Stuhrmann analysis was carried out by the fitting of the relationship^27^:

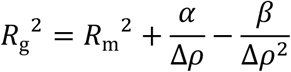

In which *R*_m_^2^ represent the *R*_g_ if it were to have a homogenous SLD. The value of a reflects the radial distribution of SLD, with values > 0 suggesting that higher contrast components are located towards the outside of the complex. The value of β is a reflection of the distance of the centre of the complex’s SLD from the complex’s centre of mass. In the case of the Stuhrmann plot of dFICD•hBiP-AMP a linear best-fit line (suggesting β ≈ 0) was a considerably better fit to the data (shown in **Fig. 2d**; R^2^ = 0.93) than the fitting of a quadratic curve (R^2^ = 0.66). Theoretical *R*_g_ values, derived from structural models, were calculated using CRYSON^58^. The symmetry of structural models was assessed through the use of AnAnaS software^59^.

### Differential Scanning Fluorimetry (DSF)

DSF experiments were performed on a CFX96 Touch Real-Time PCR Detection System (Bio-Rad) in 96-well plates (Hard-Shell, Bio-Rad) sealed with optically clear Microseal ‘B’ Adhesive Sealer (Bio-Rad). Each sample was measured in technical duplicate and in a final volume of 20 μl. Protein was used at a final concentration of 2 μM, ATP or ADP (if applicable) at 5 and 2 mM, respectively, and SYPRO Orange dye (Thermo Fisher) at a 10 × concentration in a buffer of HKM (25 mM HEPES-KOH pH 7.4, 150 mM KCl, 10 mM MgCl_2_). Solutions were briefly mixed and the plate spun at 200 *g* for 10 s before DSF measurement. Fluorescence of the SYPRO Orange dye was monitored on the FRET channel over a temperature range of 25–90 °C with 0.5 °C intervals. Background fluorescence changes were calculated and subtracted from the protein sample fluorescence data using no-protein control (NPC) wells. NPC fluorescence was unchanged by the addition of ATP or ADP. Data was then analysed in Prism 8.4 (GraphPad), with melting temperatures calculated as the global minimums of the negative first derivatives of the relative fluorescent unit (RFU) melt curves (with respect to temperature).

### Bio-layer interferometry (BLI)

AMPylated or non-AMPylated biotinylated-AviTag-haBiP^T229A-V461F^ (UK2359), was AMPylated if applicable, in vitro biotinylated (as previously described^20^), made apo and purified as detailed above in *Protein Purification.* Both proteins were confirmed as being > 95% biotinylated by streptavidin gel-shift. All BLI experiments were conducted on the FortéBio Octet RED96 System (Pall FortéBio) using in a buffer basis of HKM supplemented with 0.05% Triton X-100 (HKMTx). Streptavidin (SA)-coated biosensors (Pall FortéBio) were hydrated in HKMTx for at least 30 min at 25 °C prior to use. Experiments were conducted at 30 °C. BLI reactions were prepared in 200 μl volumes in 96-well microplates (greiner bio-one). Ligand loading was performed with biotinylated BiP-AMP:Apo at 7.5 nM and with biotinylated BiP:Apo at 5.8 nM, such that the rate of ligand loading was roughly equivalent and all tips reached a threshold of 1 nm binding signal (displacement) within 300–600 s. All ligands loaded with a range of 1.0–1.2 nm. After loading of the immobilised ligand, BiP was activated in 2 mM ATP for 200 s, followed by a 50 s baseline in HKMTx alone, before association with apo FICD variants (all bearing a catalytically inactivating His363Ala mutation and at 50 nM unless otherwise specified) in HKMTx (see schematic in **Supplementary Fig. 3d**). Note, immobilised (unmodified) BiP was previously observed to domain-dock, and remain domain-docked for extended periods of time in ATP-replete buffer, following this protocol of ATP activation^20^. The first dissociation step was initiated by the dipping of all tips into wells lacking FICD analyte (only HKMTx). The second dissociation step was induced by the dipping of the biosensor tips into HKMTx supplemented with 2 mM ATP. Experiments were conducted at a 1000 rpm shake speed and with a 5 Hz acquisition rate. Data were processed in Prism 8.4 (GraphPad).

### In vitro deAMPylation (fluorescence polarisation) assay

Measurement of deAMPylation kinetics was performed as described previously^11^ with modifications. The probe BiP^T229A-V461F^ (UK2521) modified with FAM-labelled AMP: BiP^T229A-V461F^-AMP(FAM)) was generated by pre-incubating 100 μM apo BiP^T229A-V461F^ with 5 μM GST-FICD^E234G^ (UK1479) and 110 μM ATP in HKM buffer for 5 min at 20 °C, followed by addition of 100 μM ATP-FAM [N^6^-(6-Amino)hexyl-ATP-6-FAM; Jena Bioscience] and further incubation for 19 h at 25 °C. To ensure complete BiP AMPylation 2 mM ATP was then added to the reaction which was incubated for a further 1.25 h at 25 °C. The reaction mixture was then incubated with GSH-Sepharose 4B matrix for 45 min at 4 °C in order to deplete the GST-FICD^E234G^. The BiP containing supernatant was buffered exchanged into HKM using a Zeba Spin desalting column (7K MWCO, 0.5 ml; Thermo Fisher) in order to remove the majority of free (FAM labelled) nucleotide. 2 mM ATP was added to the eluted protein and incubated for 15 min at 4 °C (to facilitate displacement of any residual FAM-labelled nucleotide derivates bound by the NBD of BiP). Pure BiP-AMP(FAM) with BiP-AMP was then obtained by gel filtration using an S75 Increase 10/300 GL column equilibrated in HKM at 4 °C. 1 mM TCEP was added to the protein fractions, which were concentrated using a 50K MWCO centrifugal filter and snap frozen. A labelling efficiency of 1.8% was estimated based on the extinction coefficient for BiP-AMP:ATP (ε_280_ 33.5 mM^-1^ cm^-1^), FAM (ε_492_ 83.0 mM^-1^ cm^-1^) and a 280/492 nm correction factor of 0.3 (Jenna Biosciences).

DeAMPylation reactions were performed in HKMTx(0.1) buffer [HKM supplemented with 0.1% (*v/v*) Triton X-100] in 384-well polysterene microplates (black, flat bottom, μCLEAR; greiner bio-one) at 30 °C in a final volume of 30 μl containing trace amounts of fluorescent BiP^T229A-V461F^-AMP(FAM) probe (10 nM), supplemented with BiP^T229A-V461F^-AMP (5 μM) and FICD proteins (0.5 μM). A well lacking FICD protein was used for baseline FP background subtraction. 10 nM ATP-FAM alone was also included as a low FP control (not shown). Under these conditions [E]_0_ was assumed to be << [S]_0_ + *K*_M_ (with [E]_0_ = 0.5 μM, [S]_0_ = 5 μM and the presumed *K*_M_ (Michaelis constant) ≥ GST-FICD *K*_M_ of 16 μM^11^) such that quasi-steady state reaction kinetics should apply with respect to the initial reaction rate. Furthermore, [S]o was considered to be sufficiently small relative to the FICD variant presumed *K*_M_ values such that, by derivation from the Michaelis-Menten equation^60^, the following relationship holds true:

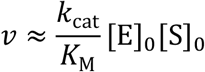

where *v* is the measured initial reaction velocity. On account of the close correspondence between the values calculated here and previously (from a Michaelis-Menten analysis of GST-FICD^11^) these assumptions are clearly valid for wild type FICD. More accurately all presented ~ *k*_cat_/*K*_M_ values are in fact equivalent to *k*_cat_/(*K*_M_ + [S]_0_).

Fluorescence polarisation of FAM (λ_ex_ = 485 nm, λ_em_ = 535 nm) was measured with an Infinite F500 plate reader (Tecan). The mFP *y*_0_ difference between the FICD^L258D^ time course and the same reaction composition pre-incubated for 5 h at 25 °C before the beginning of data collection, was interpreted as the ΔmFP equivalent to complete (5 μM) BiP-AMP deAMPylation (see **Supplementary Fig. 3e**). Fitting of the initial linear reaction phase was achieved using Prism 8.4 (GraphPad).

For direct calculation of *k*_cat_ values deAMPylation assays were conducted as above but with 10 μM FICD or FICD^L258D^ and 100 or 150 μM BiP-AMP substrate. Following subtraction of a no enzyme background from all datasets, the mFP difference for each sample (between t = 0 and the mFP plateau) was interpreted as the ΔmFP equivalent to complete BiP-AMP deAMPylation ([S]_0_).

### In vitro AMPylation

In vitro AMPylation reactions were performed in HKM buffer in a 7 μl volume. Reactions contained 10 μM ATP-FAM, 5 μM ATP-hydrolysis and substrate-binding deficient BiP^T229A-V461F^ (UK2521), 7.5 μM oxidised _S-S_FICD^A252C-H363A-C421S^ (UK2269, trap) to sequester any modified BiP [BiP-AMP(FAM)] and, unless otherwise stated, 0.5 μM FICD. Reactions were started by addition of nucleotide. Apart from in the presented time courses (**Supplementary Fig. 4b**) after a 60 min incubation at 25 °C the reactions were stopped by addition of 3 μl 3.3 × LDS sample buffer (Sigma) containing NEM (40 mM final concentration) for non-reducing SDS-PAGE or DTT (50 mM final concentration) for reducing SDS-PAGE and heated for 10 min at 70 °C. Samples were applied to a 10% SDS-PAGE gel, the FAM-label was imaged with a Chemidoc MP (Bio-Rad) using the Alexa Flour 488 dye setting. Gels were subsequently stained with Quick Coomassie (Neo Biotech).

### Mammalian Cell Culture and Lysis

The CHO-K1 *FICD^-/-^* cell line used in this study was described previously^4^. The CHO-K1 S21 *FICD^-/-^* cell line was generated by CRISPR-Cas9 knockout of both FICD alleles (as described previously^4^) into the previously described UPR reporter *CHOP::GFP* and *XBP1s::Turquoise* bearing CHO-K1 S21 cell line^61^. Cells were cultured as in^20^. Where indicated, cells were treated for 3 h with cycloheximide (Sigma) by exchanging the culture medium with pre-warmed (37 °C) medium supplemented with cycloheximide at 100 μg/ml. Cell lysates were obtained and analysed as in^20^ but with a HG lysis buffer consisting of 20 mM HEPES-KOH pH 7.4, 150 mM NaCl, 2 mM MgCl_2_, 33 mM D-glucose, 10% (*v/v*) glycerol, 1% (*v/v*) Triton X-100 and protease inhibitors (2 mM phenylmethylsulphonyl fluoride (PMSF), 4 μg/ml pepstatin, 4 μg/ml leupeptin, 8 μg/ml aprotinin) with 100 U/ml hexokinase (from *Saccharomyces cerevisiae* Type F-300; Sigma).

### Immunoblot (IB) analysis

After separation by SDS-PAGE or native-PAGE (as previously described^20^) proteins were transferred onto PVDF membranes. The membranes were blocked with 5% *(w/v)* dried skimmed milk in TBS (25 mM Tris-HCl pH 7.5, 150 mM NaCl) and incubated with primary antibodies followed by IRDye fluorescently labelled secondary antibodies (LI-COR). The membranes were scanned with an Odyssey near-infrared imager (LI-COR). Primary antibodies and antisera against hamster BiP [chicken anti-BiP^62^], eIF2α [mouse anti-eIF2α^63^] and FICD [chicken anti-FICD^4^] were used.

### Flow cytometry

FICD over-expression-dependent induction of unfolded protein response signalling was analysed by transient transfection of CHO-K1 S21 *FICD^-/-^* UPR reporter cell lines with plasmid DNA encoding the complete FICD coding sequence (with mutations as indicated) and mCherry as a transfection marker, using Lipofectamine LTX (Thermo Fisher) as described previously^4^. 0.5 μg DNA was used to transfect cells growing in 12-well plates. 40 h after transfection the cells were washed with PBS and collected in PBS containing 4 mM EDTA, and single live-cell fluorescent signals (20,000 collected per sample) were analysed by dual-channel flow cytometry with an LSRFortessa cell analyser (BD Biosciences). Turquoise and mCherry fluorescence was detected using a 405 nm excitation laser with a 450/50 nm emission filter and a 561 nm excitation laser with a 610/20 nm emission filter, respectively. Data were processed using FlowJo and the extracted population parameters were plotted in Prism 8.4 (GraphPad).

## Data availability

The deAMPylation complex crystal structures of monomeric FICD and AMPylated BiP have been deposited in the Protein Data Bank (PDB) with the following accession codes: 7B7Z (State 1), 7B80 (State 2). Raw SANS data is available from doi:10.5291/ILL-DATA.8-03-963.

## Acknowledgements

We thank the Huntington lab for access to the Octet machine and the CIMR flow cytometry core facility team (Reiner Schulte, Chiara Cossetti and Gabriela Grondys-Kotarba). This work was supported by Wellcome Trust Principal Research Fellowship to D.R. (Wellcome 200848/Z/16/Z), and a Wellcome Trust Strategic Award to the Cambridge Institute for Medical Research (Wellcome 100140). We are grateful to the Diamond Light Source for X-ray beamtime (proposal MX-21426) and the staff of beamline I04-1 for assistance with data collection; and to the ILL for neutron beamtime as part of proposal 8-03-963 with particular thanks to Anne Martel for her assistance. For advice pertaining to the use of PEPSI-SANS software we thank Sergei Grudinin. We are indebted to Yahui Yan for the gift of FICD’s TPR domain-expressing plasmid and to Cláudia Rato da Silva for advice and guidance on the in vivo experiments. We also thank Yahui Yan, Alisa F. Zyryanova and Lisa Neidhardt for comments on the manuscript.

## Author contributions

L.A.P. led and conceived the project, designed and conducted the experiments, analysed and interpreted all the data, purified and crystallised proteins, collected, analysed and interpreted the X-ray diffraction and neutron scattering data, and wrote the manuscript. S.P.^1^ conducted the FP assays, purified proteins. N.Z. and S.P.^2^ helped to collect and process scattering data. J.M.D. and M.H. expressed the deuterated proteins. All authors contributed to revising the article. D.R. conceived and oversaw the project, interpreted the data, and wrote the manuscript.

## Conflict of interests

The authors declare no conflict of interests.

## Notes

### Competing Interest Statement

The authors have declared no competing interest.

