## Supplementary Figures, Legends and Tables for "Structures of a deAMPylation complex rationalise the switch between antagonistic catalytic activities of FICD (14/96/109)"

### Supplementary Fig. 1

**a**

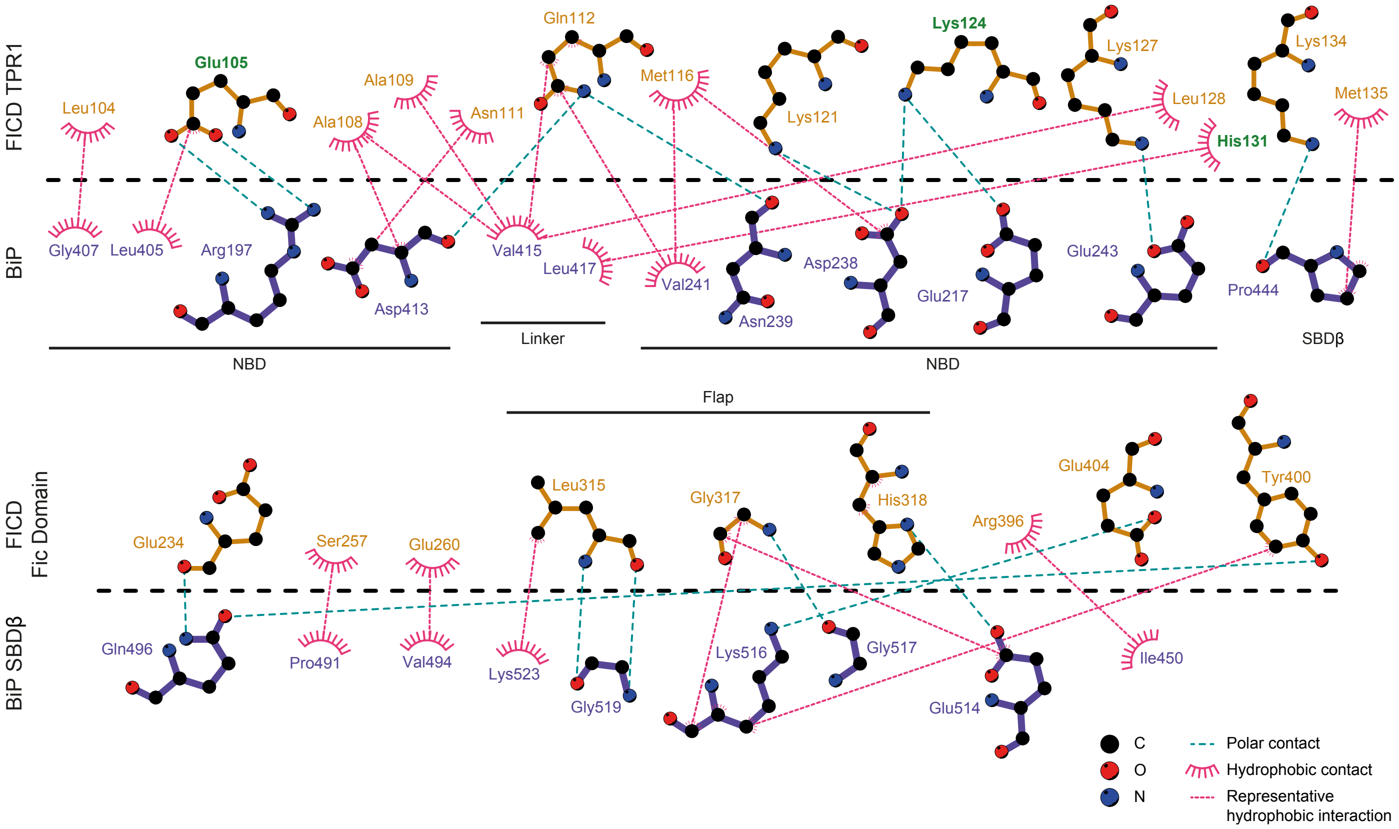


**b**

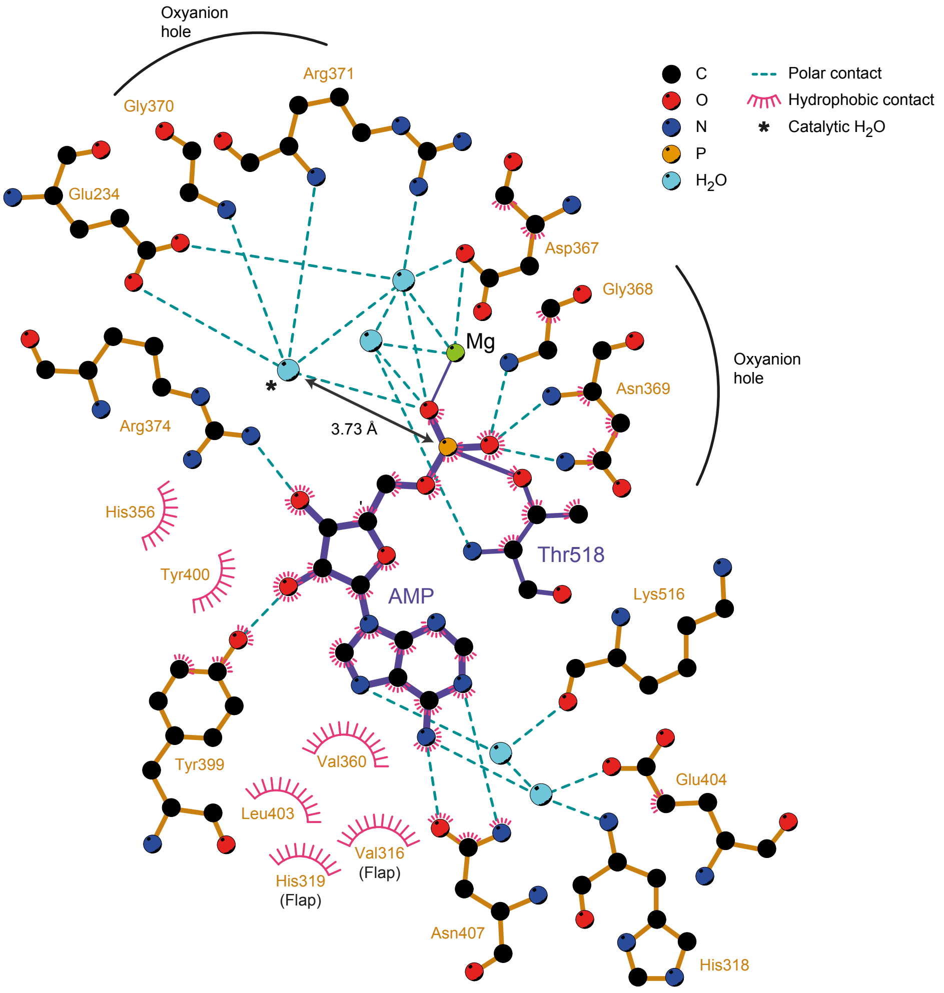

**Supplementary Fig. 1: Schematised view of FICD•BiP-AMP intermolecular contacts. a,** All polar (hydrogen bonds and salt bridges) and hydrophobic contacts between FICD and BiP are illustrated. The (sub)domain origin of the interacting residue is also annotated. Residues mutated in the study are labelled in green. **b,** Intermolecular contacts between BiP's Thr518-AMP, $Mg^{2+}$ and FICD's catalytic domain, as shown in **Fig. 1d**, are depicted. Various Fic domain features and the distance between the catalytic water* and the AMP phosphor atom and are also annotated. Note, the tight binding of the adenosine and α-phosphate along with Arg374 coordination of the ribose 3'-OH would prohibit the intramolecular nucleophilic attack and cyclisation required for an anchimeric-assisted mechanism of BiP deAMPylation (an alternative mechanism capable of generating the experimentally-observed deAMPylation products, unmodified BiP and AMP[11]). Likewise, the lack of base (required for proton abstraction) in the vicinity of BiP's Thr518 Cα is inimical to an E1cB-type elimination-based deAMPylation reaction.

### Supplementary Fig. 2

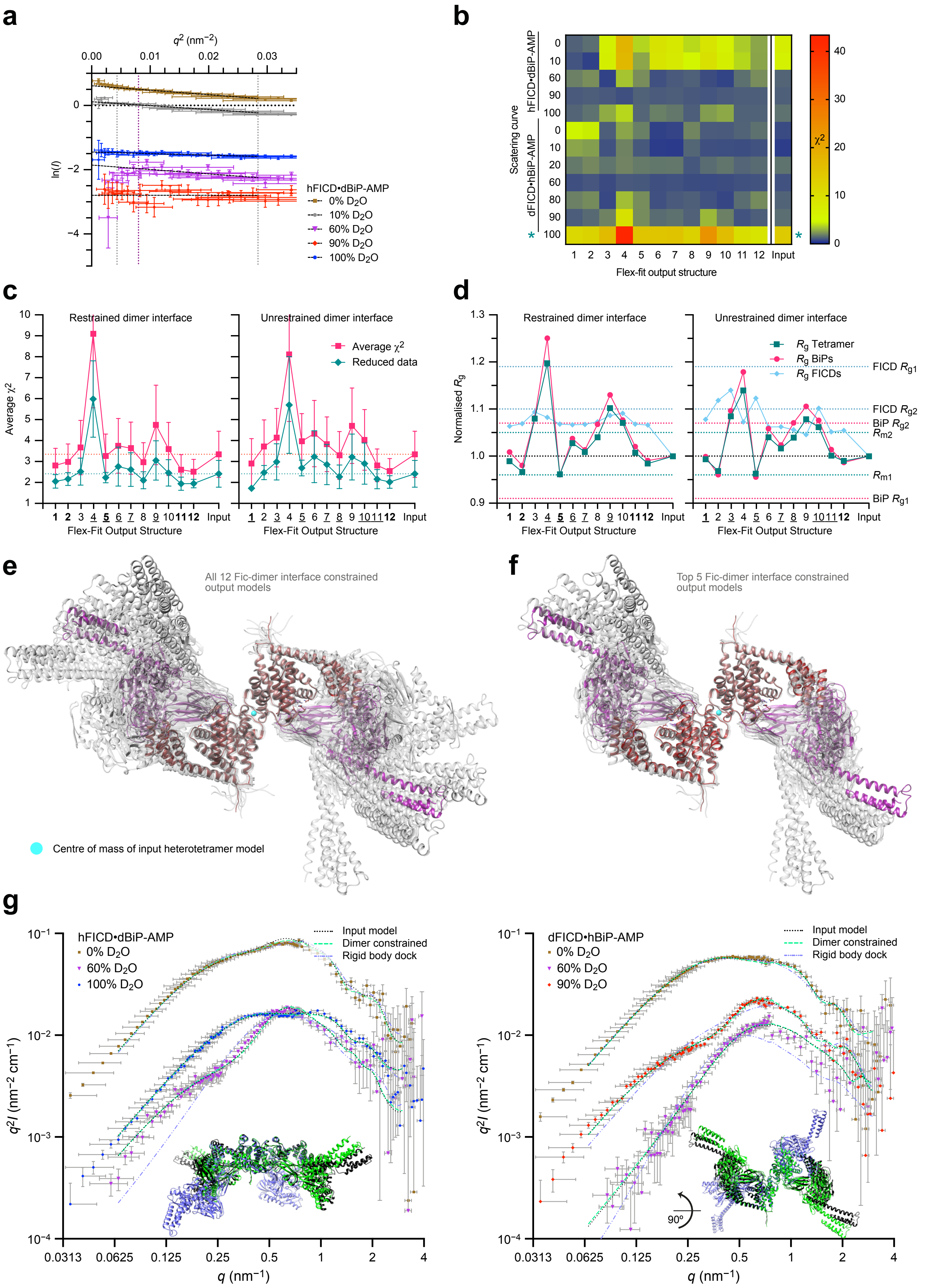

**Supplementary Fig. 2: SANS data analysis. a,** Guinier plot of non-deuterated FICD with partially deuterated AMPylated BiP. **b,** Heat-map of the $\chi^2$ goodness of fit of the theoretical scattering curve of each flex-fit model against all observed experimental scattering datasets. The major NW to SE diagonal illustrates the optimised $\chi^2$ of each flex-fit model from its progenitor dataset. **c,** Comparison of the mean $\chi^2$ for each model derived from analysis of the goodness of fit to all scattering datasets (generated from both oppositely labelled complexes). The 'reduced data' average $\chi^2$ (green) is derived from fitting to all data excluding the anomalous scattering observed for dFICD•hBiP-AMP in 100% $D_2O$ (***b**). Error bars are SEM. **d,** Comparison of Stuhrmann analysis derived $R_g$s (horizontal dotted lines) with the calculated $R_g$s of the input and output structures. In **c** and **d** output structures highlighted in bold have $\chi^2$ SDs (for reduced data) which are less than and significantly different to the input model's $\chi^2$ SD ($P < 0.05$ by F-test); symmetrical output structures are underlined. **e,** Superposition of all 12 flex-fit output structures (with FICD dimer interface constrained) relative to the input heterotetramer model (red FICD dimer, purple BiP-AMPs). **f,** As in **e** but only displaying the top 5 flex-fit structures with significantly reduced $\chi^2$ SDs. **g,** Kratky plots of representative scattering curves highlighting the relative fits of the input, dimer constrained best-fit and a poorer fitting rigid-body docking models. Inset, colour-matched structures aligned by the FICD dimer, shown in orthogonal views. Note, the scattering intensity profiles are consistent with FICD•BiP-AMP being a folded protein complex. See **Supplementary Movie 1**.

### Supplementary Fig. 3

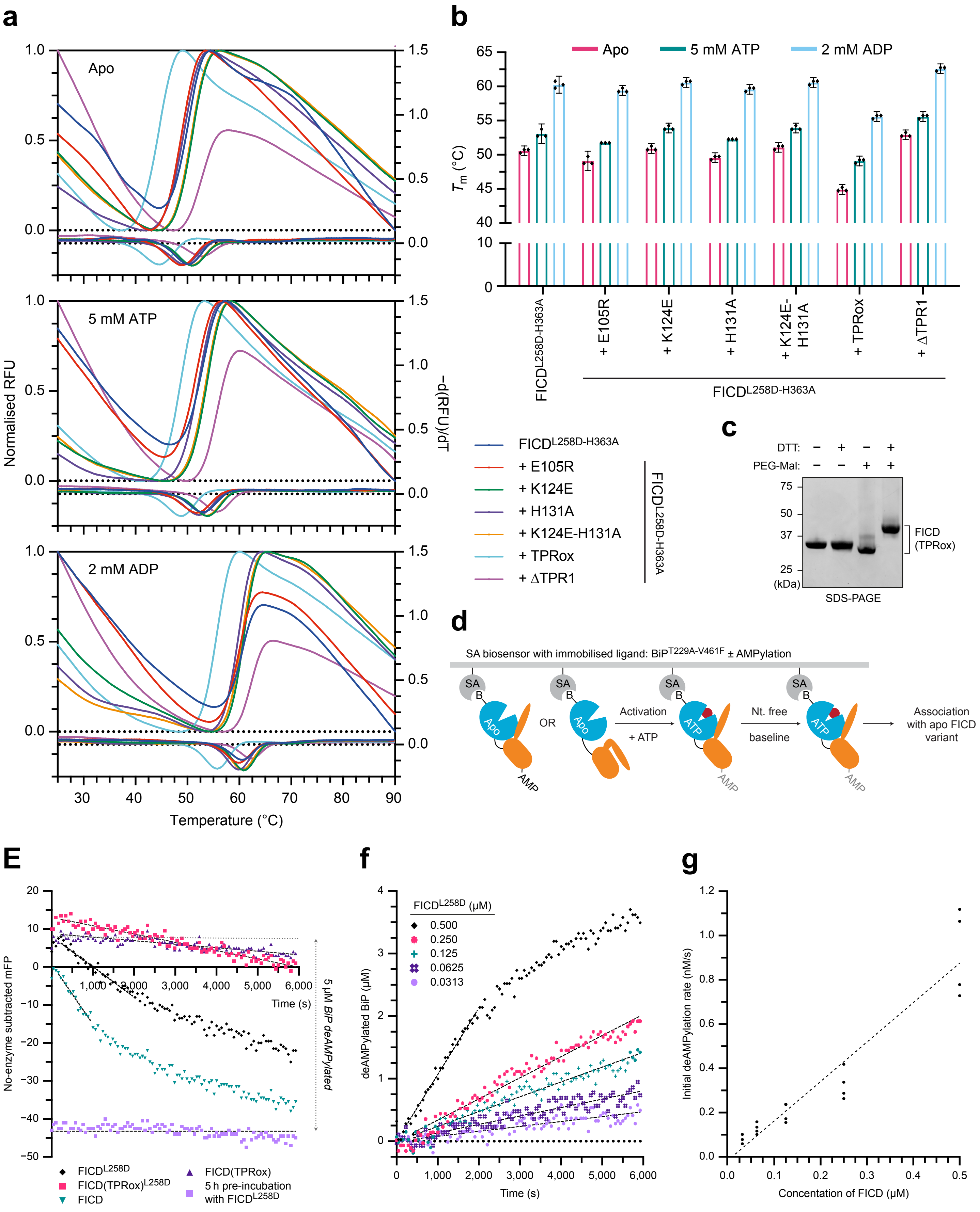

**Supplementary Fig. 3: Biophysical analysis of FICD mutants and in vitro deAMPylation assay. a,** Representative normalised DSF melt curves (top curves) with corresponding negative first derivatives (bottom curves). **b,** The derived protein melting temperatures ($T_m$, mean ± 95% CI) derived from n = 3 independent DSF experiments each conducted in technical duplicate. Note, all FICD variants (like $FICD^{L258D-H363A}$) are stabilised by nucleotide binding. **c,** PEG 2000 maleimide-based electrophoretic mobility assay analysis of the oxidation status of monomeric $FICD^{L258D-H363A}$(TPRox), demonstrating almost complete disulphide-stapling of the FICD's TPR domain to its linker helix. **d,** Schematised model of the BLI protocol (for immobilised BiP ± AMP) preceding the association and dissociation phases shown in **Fig. 3**. **e,** The FP curves from which **Fig. 3c** was derived. The difference (in mFP units) at t = 0 between the $FICD^{L258D}$ deAMPylation time course and the pre-incubated and fully deAMPylated reaction was taken to represent complete substrate deAMPylation (5 µM BiP-AMP). Fits of the linear enzyme velocities are overlaid. **f,** The FP-converted time course of BiP-AMP(FAM) deAMPylation with different concentrations of $FICD^{L258D}$. **g,** Quantification of the assay represented in **f**, from n = 4 independent experiments — demonstrating the minimum linear dynamic range of the assay. The dashed line illustrates the unconstrained best-fit linear relationship.

### Supplementary Fig. 4

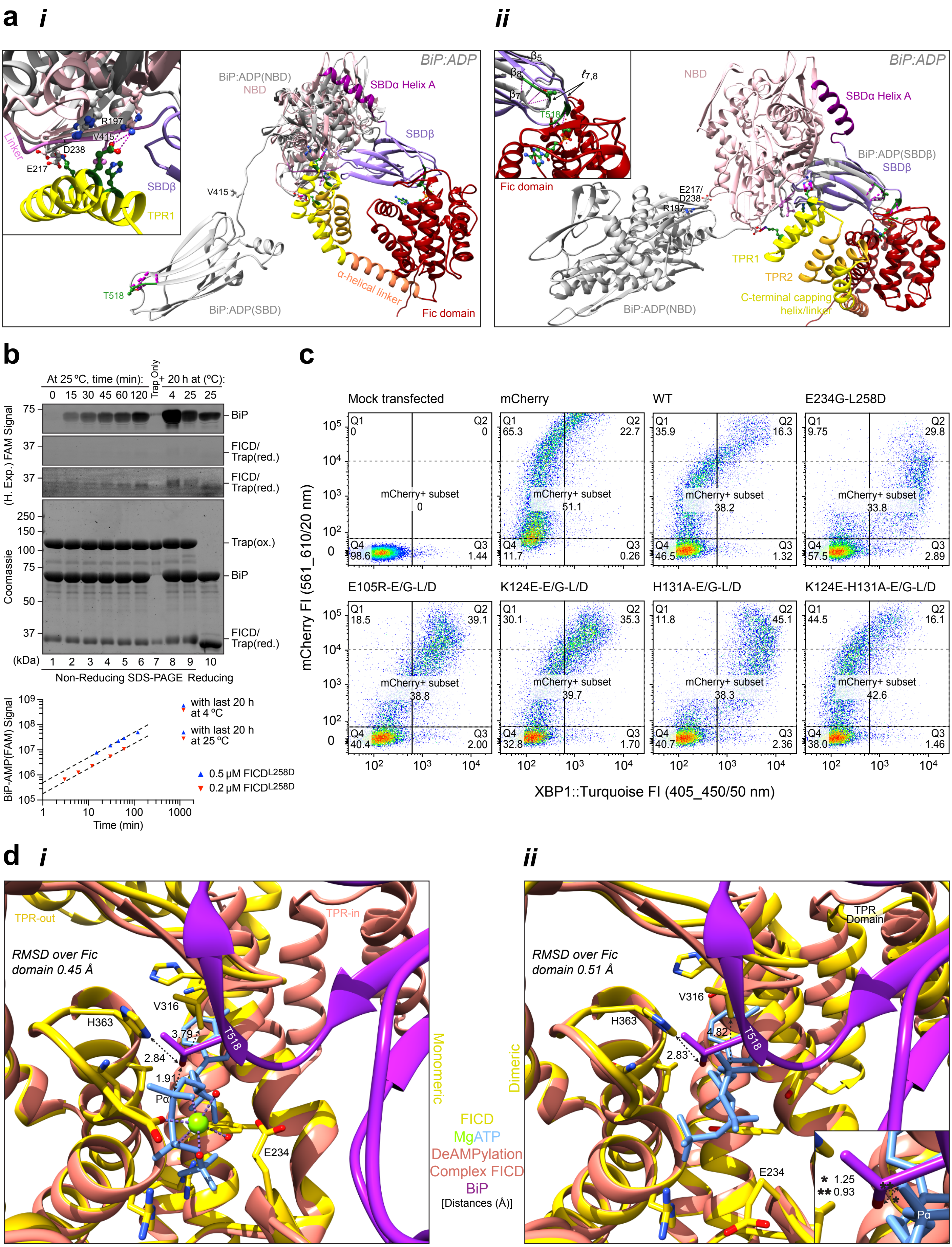

**Supplementary Fig. 4: FICD recognises the ATP-state of BiP. a,** The deAMPylation complex (coloured as in **Fig. 1b** with selected BiP interaction partners labelled) is aligned via its NBD (***i***) or SBDβ (***ii***) with an ADP-state BiP (PDB 7A4U; grey). Inset (***i***), a closeup view of FICD(TPR1)-BiP(NBD) contacts. (***ii***), the intermolecular β-sheet region of $\ell_{7,8}$ (green) is shortened in BiP:ADP. Inset, disposition of Thr518 is highlighted (pink lines; H-bonds). See **Supplementary Movie 2**. **b,** Representative $FICD^{L258D}$ (0.5 μM) BiP-AMPylation time course (experimental design as in **Fig. 4b**) with quantification of two independent experiments (dashed lines; directly proportional fits). **c,** FACS dot-plot. Cells with mCherry+ signal $\leq 10^4$ were gated, eliminating the distorting effect of high FICD expression (marked by the mCherry) on the XBP1::Turquoise signal, and further analysed in **Fig. 4d**. **d,** (***i***) Modelled pre-AMPylation complex illustrating the ability of monomeric FICD, bound to MgATP (PDB 6I7K; yellow with purple nucleotide), to accommodate BiP's Thr518 in a catalytically-competent conformation. Derived from alignment with FICD from the deAMPylation complex (orange; BiP in purple). Oγ of BiP's Thr518 (restored by removal of the AMP) is in-line with the $P\alpha$-$O^{3\alpha}$-phosphoanhydride bond and can be deprotonated by His363. Though not modelled here, flexibility in the Fic flap, FICD's Val316 and BiP($\ell_{7,8}$) likely permit Pα and Oγ(Thr518) to attain a distance consistent with an initial substrate engagement state. (***ii***) As in ***(i)*** but with alignment of dimeric FICD bound to ATP in a catalytically-incompetent mode (PDB 6I7G). Note the severe clash between Thr518 and the ATP α-phosphate (*inset).

### Supplementary Fig. 5

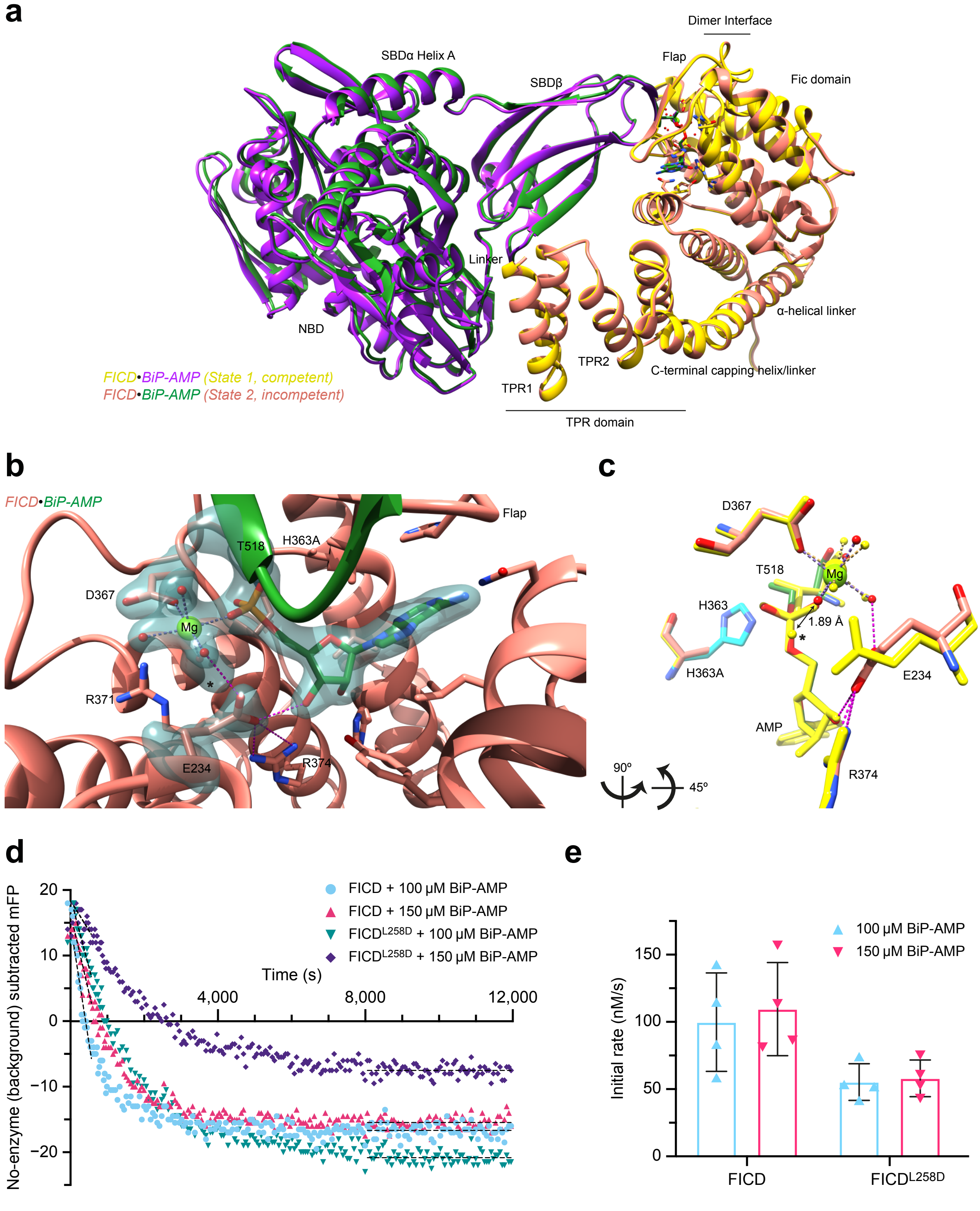

**Supplementary Fig. 5: A second deAMPylation complex crystal structure captures a non-catalytically competent state of monomeric FICD. a,** State 1 and state 2 heterodimeric deAMPylation complexes superposed. See **Supplementary Movie 1**. **b,** State 2 deAMPylation complex focus on the active site, with regions of particular interest additionally overlaid with an unbiased polder-omit electron density map, contoured at 6σ. See **Supplementary Movie 3**. In **a** and **b** structures are depicted in the same view as the state 1 complex shown in **Fig. 1b** and **Fig. 1d**, respectively. **c,** The same reduced active site view as shown in **Fig. 5a**, with the polder-omit map removed for clarity. In **b** and **c** interactions formed by state 2's Glu234 are shown with pink-dashed lines. **d,** Background drift–subtracted FP deAMPylation time course, the basis of the panel displayed in Fig 5b. Linear best-fits are overlaid illustrating the initial reaction progress and final plateau value, the ΔFP between $y_0$ and $y_\infty$ was taken to represent $[\text{BiP-AMP}]_0$. **e,** Quantification of the initial deAMPylation rates (mean ± SD) with either 100 or 150 µM BiP-AMP substrate at t = 0. Results are presented from n = 4 independent experiments.

### Supplementary Fig. 6

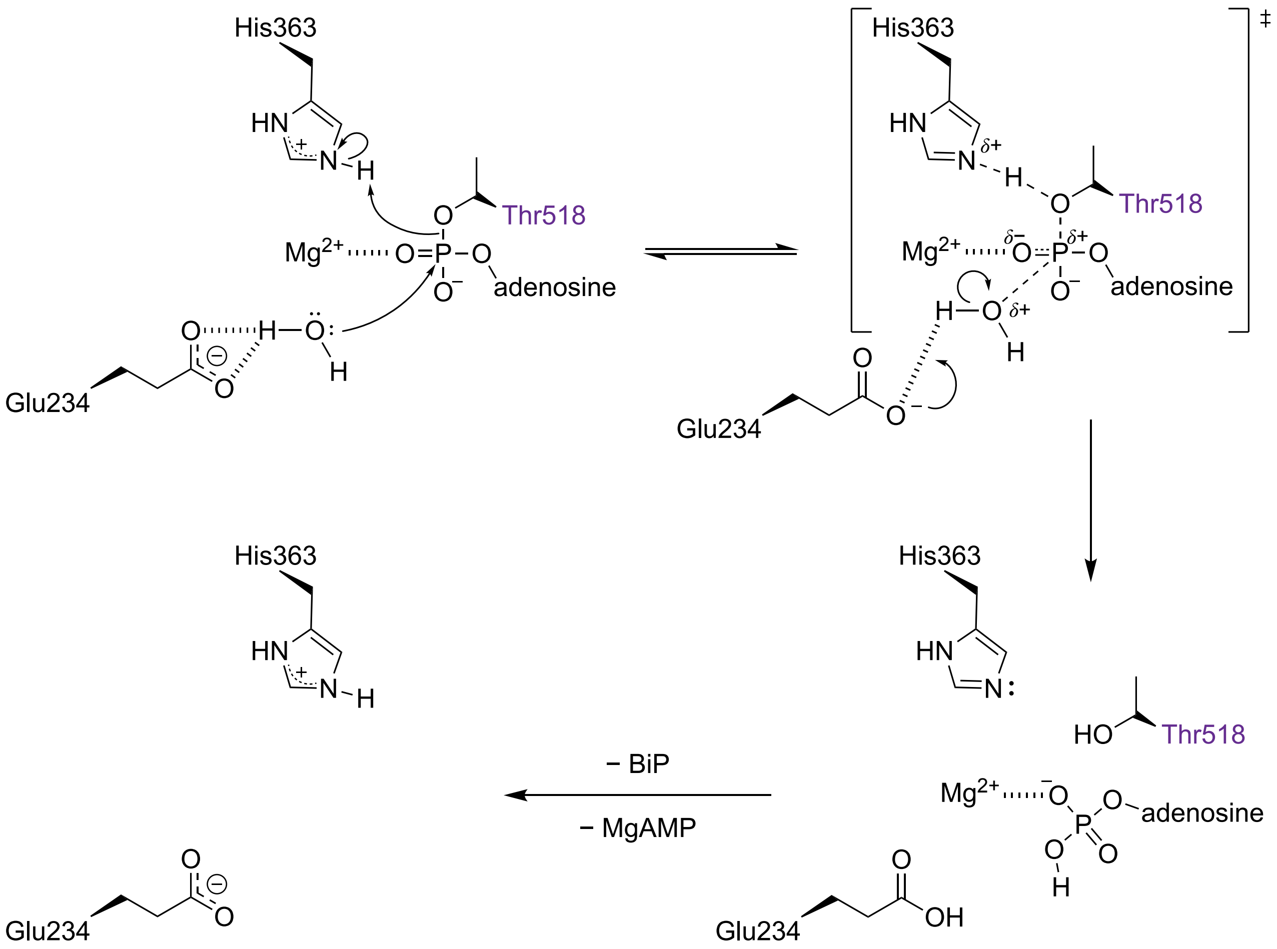

**Supplementary Fig. 6: Proposed hydrolytic BiP deAMPylation mechanism.** FICD's Glu234 activates and aligns a catalytic water molecule for in-line nucleophilic attack into the backside of AMPylated BiP's Pα-Oγ(Thr518) phosphodiester bond. The α-phosphate group coordination by $Mg^{2+}$, and localisation within FICD's electron withdrawing oxyanion hold (**Supplementary Fig. 1b**), stabilises the position of Pα and increases its electrophilicity. His363 can exist in either a protonated or deprotonated state. The former is required for deAMPylation and is shown. The reaction likely proceeds via a nucleophilic $S_N2$-type pathway with concerted protonation of BiP's Thr518 alkoxide leaving group (catalysed by FICD's His363 acting as a general acid). A potential role for FICD's Glu234 acting as a catalytic (but not general) base, accepting a proton from the nucleophilic water at a late stage of the reaction after formation of the pentacoordinate transition state (‡), is shown. Glu234 acting as a proton trap is consistent with its interaction with Fic motif Arg371 and Arg374 (not shown; **Fig. 1e**), which will depress its $pK_a$. The schematised hydrolytic reaction generates BiP (with an unmodified Thr518) and AMP. Following product release Glu234 and His363 could facilely exchange protons with the solvent to regenerate the original FICD active site. Polar interactions are denoted with hashed lines, dashed lines represent partial covalent bonds and partial charges are indicated by δ. BiP's Thr518 residue is annotated in purple.

| | Input model | hFICD•dBiP-AMP | dFICD•hBiP-AMP | Best-fit (dimer constrained) | Best-fit (dimer unconstrained) |
|---|---|---|---|---|---|
| **$M_W$ (kDa)** | 220 | 220 ± 10 | 250 ± 30 | | |
| **$R_m$ [$R_g$ of complex] (Å)** | 60.1 | 58 ± 19 (43 to 70) | 63 ± 14 (58 to 68) | 57.8 | 59.7 |
| **$R_g$ of FICDs (Å)** | 34.5 | 41 (17 to 55) | 38 (14 to 52) | 36.9 | 37.2 |
| **$R_g$ of BiPs (Å)** | 69.0 | 63 (52 to 73) | 74 (69 to 79) | 66.3 | 69.0 |
| **$\chi^2$ (reduced data)** | 2.4 ± 2 | | | 2.4 ± 0.8 | 1.7 ± 0.4 |

**Supplementary Table 1. SANS data summary.** Biophysical parameters derived from forward scattering (molecular weight) and Stuhrmann analysis of the contrast variation SANS data over the low-*q* (Guinier) region are shown. The molecular weight ($M_W$) is estimated by comparison of experimental and theoretical *I*(0)/c values (the latter assuming an amino acid composition of a 1:1 complex of FICD and BiP-AMP and taking into account buffer component contributions). The mean $M_W$ ± SD is calculated across all curves excluding the 60% $D_2O$ datasets, which are close to the contrast match points for both partially deuterated complexes. Radii of gyration parameters are best fit or interpolated values ± SE (and/or 95% CI) from the Stuhrmann curve fittings (**Fig. 2d**). Parameters from the best flex-fit heterotetramer models are shown with their theoretical $R_g$ values and overall $\chi^2$ goodness of fit (mean ± SD) against the reduced scattering dataset (**Fig. 2e** and **Supplementary Fig. 2c**). The $\chi^2$ of the input model, against the reduced scattering dataset, is also shown for reference (**Fig. 2a** and **Supplementary Fig. 2c**).

| ID | Plasmid name | Description | Encoded protein | Figure | Purification | PMID |
|---|---|---|---|---|---|---|
| **UK1983** | yUlp1(402-621)-StrepII_pET24a | Bacterial expression of yeast Ulp1 protease with a C-terminal StrepII tag | Ulp1-StrepII | | | 31531998 |
| **UK1479** | hsHYPE_45-458_E234G_pGV67 | Bacterial expression of hyperactive human GST-TEV-HYPE_45-458 | GST-FICD$^{E234G}$ | | | 26673894 |
| **UK2090** | haBiP_27–549_T229A_V461F_pQE30_pSmt3 | Bacterial expression of ATPase and substrate binding deficient, lid truncated BiP (N-terminal H6-Smt3 fusion) | Lid truncated BiP$^{T229A\text{-}V461F}$ | 1 | RQ, S75pg | 33295873 |
| **UK2093** | hsHYPE_104–445_L258D_H363A_pSmt3_pET28b | Bacterial expression of inactive and monomeric human H363A mutant of His6-Smt3-HYPE(104–445) | FICD$^{L258D\text{-}H363A}$ | 1, 3A–B, 4A | RQ, S75pg | 31531998 |
| **UK2521** | haBiP_27–635_T229A_V461F_pSmt3_pET28b | Bacterial expression of FL ATPase and substrate binding deficient His6-Smt3-BiP(27–635) | BiP$^{T229A\text{-}V461F}$ | 2 | RQ, S75pg | |
| **UK1954** | hsHYPE_104–445_H363A_pSmt3_pET28b | Bacterial expression of inactive human H363A mutant of His6-Smt3-HYPE(104–445) | FICD$^{H363A}$ | 2, 3A–B, 4A | RQ, S200pg | |
| **UK1801** | EcBirA_WT_pGEX_TEV | Bacterial expression of fastidious E. coli BirA biotin ligase (R118 intact) | GST-BirA | | | |
| **UK2359** | haBiP_27–635_T229A_V461F_pQE30_Smt3_Avi | FL hamster BiP ATPase dead, substrate binding deficient with N-terminal H6-SUMO-AviTag and small GS linker | Biotinylated-BiP$^{T229A\text{-}V461F}$ | 2 | MQ | |
| **UK2054** | hsHYPE_138-445_H363A_pSmt3_pET28b | Bacterial expression of H363A ΔTPR1 mutant of human His6-Smt3-HYPE(138-445) | FICD(ΔTPR1)$^{H363A}$ | 3A | MQ, S200Incr | |
| **UK2051** | hsFICD_104–186_pSUMO(M3) | Bacterial expression of human FICD/HYPE TPR domain, residues 104–186 | TPR Domain | 3A | RQ, S75Incr | |
| **UK2607** | hsHYPE_104–445_E105R_L258D_H363A_pSmt3_pET28b | Bacterial expression of inactive human L258D H363A mutant of His6-Smt3-HYPE_104–445 with TPR mutation | FICD$^{E105R\text{-}L258D\text{-}H363A}$ | 3B(i) | RQ, S75Incr | |
| **UK2617** | hsHYPE_104–445_K124E_H131A_L258D_H363A_pSmt3_pET28b | Bacterial expression of inactive human L258D H363A mutant of His6-Smt3-HYPE_104–445 with 2x TPR mutation | FICD$^{K124E\text{-}L258D\text{-}H363A}$ | 3B(i) | RQ, S75Incr | |
| **UK2610** | hsHYPE_104–445_H131A_L258D_H363A_pSmt3_pET28b | Bacterial expression of inactive human L258D H363A mutant of His6-Smt3-HYPE_104–445 with TPR mutation | FICD$^{H131A\text{-}L258D\text{-}H363A}$ | 3B(i) | RQ, S75Incr | |
| **UK2617** | hsHYPE_104–445_K124E_H131A_L258D_H363A_pSmt3_pET28b | Bacterial expression of inactive human L258D H363A mutant of His6-Smt3-HYPE_104–445 with 2x TPR mutation | FICD$^{K124E\text{-}H131A\text{-}L258D\text{-}H363A}$ | 3B(i) | RQ, S75Incr | |
| **UK2612** | hsHYPE_104_445_D160C_T183C_L258D_H363A_C421S_pSmt3_pET28b | Bacterial expression of inactive human L258D H363A mutant of His6-Smt3-HYPE_104–445 with TPR stapleable cysteines | FICD$^{L258D\text{-}H363A}$(TPRox) [$_{s\text{-}s}$ FICD$^{D160C\text{-}T183C\text{-}L258D\text{-}H363A\text{-}C421S}$] | 3B(i) | CQ, RQ, S75Incr | |
| **UK2579** | hsHYPE_104–445_E105R_H363A_pSmt3_pET28b | Bacterial expression of inactive human H363A mutant of His6-Smt3-HYPE_104–445 with TPR mutation | FICD$^{E105R\text{-}H363A}$ | 3B(ii) | RQ, S200Incr | |
| **UK2582** | hsHYPE_104–445_K124E_H363A_pSmt3_pET28b | Bacterial expression of inactive human H363A mutant of His6-Smt3-HYPE_104–445 with TPR mutation | FICD$^{K124E\text{-}H363A}$ | 3B(ii) | RQ, S200Incr | |
| **UK2583** | hsHYPE_104–445_H131A_H363A_pSmt3_pET28b | Bacterial expression of inactive human H363A mutant of His6-Smt3-HYPE_104–445 with TPR mutation | FICD$^{H131A\text{-}H363A}$ | 3B(ii) | RQ, S200Incr | |

| | | | | | | |
|---|---|---|---|---|---|---|
| **UK2675** | hsHYPE_104–445_K124E_H131A_H363A_pSmt3_pET28b | Bacterial expression of inactive human H363A mutant of His6-Smt3-HYPE_104–445 with 2 TPR mutations | FICD$^{K124E\text{-}H131A\text{-}H363A}$ | 3B(ii) | RQ, S200Incr | |
| **UK2296** | hsHYPE_104_445_D160C_T183C_H363A_C421S_pSmt3_pET28b | Dimeric FICD(H363A) made cysteine free apart from disulphide stapleable TPRs | FICD$^{H363A}$(TPRox) [$_{s\text{-}s}$FICD$^{D160C\text{-}T183C\text{-}H363A\text{-}C421S}$] | 3B(ii) | CQ, RQ, S200Incr | |
| **UK2091** | hsHYPE_104–445_L258D_pSmt3_pET28b | Bacterial expression of monomeric His6-Smt3-HYPE_104–445. Enzymatically active. | FICD$^{L258D}$ | 3C, 4B, 5 | RQ, S75pg | 31531998 |
| **UK2760** | hsHYPE_104–445_K124E_L258D_pSmt3_pET28b | Monomeric His6-Smt3-HYPE_104–445 TPR mutant, bacterial expression. Enzymatically active. | FICD$^{K124E\text{-}L258D}$ | 3C, 4B | RQ, S75Incr | |
| **UK2761** | hsHYPE_104–445_H131A_L258D_pSmt3_pET28b | Monomeric His6-Smt3-HYPE_104–445 TPR mutant, bacterial expression. Enzymatically active. | FICD$^{H131A\text{-}L258D}$ | 3C, 4B | RQ, S75Incr | |
| **UK2759** | hsHYPE_104–445_K124E_H131A_L258D_pSmt3_pET28b | Monomeric His6-Smt3-HYPE_104–445 TPR double mutant, bacterial expression. Enzymatically active. | FICD$^{K124E\text{-}H131A\text{-}L258D}$ | 3C, 4B | RQ, S75Incr | |
| **UK2762** | hsHYPE_138–445_L258D_pSmt3_pET28b | Monomeric ΔTPR1 His6-Smt3-HYPE(138–445), bacterial expression. Enzymatically active. | FICD(ΔTPR1)$^{L258D}$ | 3C, 4B | RQ, S75Incr | |
| **UK2052** | hsHYPE_104–445_pSmt3_pET28b | Bacterial expression of wild type His6-Smt3-HYPE(104–445) | FICD | 3C, 4B, 5 | RQ, S200pg | 31531998 |
| **UK2763** | hsHYPE_104_445_D160C_T183C_C421S_pSmt3_pET28b | FICD bacterial expression. TPR stapleable. Enzymatically active. | FICD(TPRox) [$_{s\text{-}s}$FICD$^{D160C\text{-}T183C\text{-}C421S}$] | 3C, 4B | CQ, RQ, S75Incr | |
| **UK2764** | hsHYPE_104_445_D160C_T183C_L258D_C421S_ pSmt3_pET28b: | Monomeric FICD bacterial expression. TPR stapleable. Enzymatically active. | FICD$^{L258D}$(TPRox) [$_{s\text{-}s}$FICD$^{D160C\text{-}T183C\text{-}L258D\text{-}C421S}$] | 3C, 4B | CQ, RQ, S75Incr | |
| **UK2269** | hsHYPE_104–445_A252C_H363A_C421S_pSmt3_pET28b | Catalytically dead and constitutively dimeric (disulphide stapleable dimer interface) FICD. Trap for BiP-AMP. | Trap [$_{s\text{-}s}$FICD$^{A252C\text{-}H363A\text{-}C421S}$] | 4B | RQ, S200pg | 31531998 |
| **UK1314** | pCEFL_mCherry_3XFLAG_C | Mammalian expression of C-terminally 3xFLAG. Neomycin-resistance replaced by mCherry (under SV40 promoter control) | mCherry | 4C, 4D | | 25858979 |
| **UK1397** | hsHYPE_WT_pCEFL_mCherry | FL CDS of WT HYPE in pCEFL marked with mCherry | FICD | 4C, 4D | | 26673894 |
| **UK2139** | hsHYPE_E234G_L258D_pCEFL_mCherry | Mammalian expression of full-length human FICD with E234G and L258D mutations in pCEFL marked with mCherry | FICD$^{E234G\text{-}L258D}$ | 4C, 4D | | 31531998 |
| **UK2676** | hsHYPE_E105R_E234G_L258D_pCEFL_mCherry | Mammalian expression of FL CDS of L258D E234G HYPE plus TPR mutation in pCEFL marked with mCherry | FICD$^{E105R\text{-}E234G\text{-}L258D}$ | 4C, 4D | | |
| **UK2677** | hsHYPE_K124E_E234G_L258D_pCEFL_mCherry | Mammalian expression of FL CDS of L258D E234G HYPE plus TPR mutation in pCEFL marked with mCherry | FICD$^{K124E\text{-}E234G\text{-}L258D}$ | 4C, 4D | | |
| **UK2678** | hsHYPE_H131A_E234G_L258D_pCEFL_mCherry | Mammalian expression of FL CDS of L258D E234G HYPE plus TPR mutation in pCEFL marked with mCherry | FICD$^{H131A\text{-}E234G\text{-}L258D}$ | 4C, 4D | | |
| **UK2679** | hsHYPE_K124E_H131A_E234G_L258D_pCEFL_mCherry | Mammalian expression of FL CDS of L258D E234G HYPE plus 2 TPR mutations in pCEFL marked with mCherry | FICD$^{K124E\text{-}H131A\text{-}E234G\text{-}L258D}$ | 4C, 4D | | |

**Supplementary Table 2. List of plasmids used in the study.** *ID* denotes the unique lab identification (UK) number of each plasmid. *Purification* contains information pertaining to any FPLC columns used in the purification of the protein (following strep-tactin, GSH-Sepharose, Ni-NTA affinity chromatography, and on bead cleavage and elution, as appropriate). CQ, HiTrap 5 ml capto Q; RQ, RESOURCE Q 6 ml; MQ, Mono Q 5/50 GL; S75pg, HiLoad 16/60 Superdex 75 prep grade; S200pg, HiLoad 16/60 Superdex 200 prep grade; S75Incr, S75 Increase 10/300 GL; S200Incr, S200 Increase 10/300 GL. *PMID* specifies references to previous publications in which the relevant plasmid has been used, if applicable. FL; full-length.

#### Description of Additional Supplementary Files

**Supplementary Movie 1: Global architecture of the eukaryotic deAMPylation complex**. The similarity of the state 1 and state 2 heterodimeric deAMPylation crystal structures of FICD•BiP-AMP is apparent, as is the similarity of the bound BiP-AMP molecule with the isolated structure of BiP:ATP. In the frames that follow we have modelled a heterotetrameric deAMPylation complex by imposing the symmetry of dimeric FICD, and the lid structure of full-length BiP:ATP. Note, the model is compatible with the membrane topology of FICD and is supported by SANS data. Morphs and alignments with the best-fit solution structure (derived from flex-fit SANS analysis with no constraints placed on the FICD dimer interface) appear next. These are suggestive of the existence of increased deAMPylation complex flexibility in solution, relative to the situation in crystallo, especially in the disposition of the BiP SBDα and NBD and to a lesser extent in the FICD TPR domain.

**Supplementary Movie 2: FICD recognises the Hsp70 ATP-state of BiP**. Alignment of BiP:ADP with the NBD of BiP-AMP from the heterodimeric deAMPylation complex are depicted. Selected interacting residue pairs in the deAMPylation complex are shown and labelled with hydrogen bonds (pink dashed lines) and representative hydrophobic contacts (blue dashed lines) where appropriate. The same BiP residues in BiP:ADP are also illustrated, highlighting the disruption of NBD-linker interface that occurs in the BiP ADP-state and the resulting incompatibility of BiP:ADP interaction with FICD(TPR). The frames that follow superimpose BiP:ADP and BiP-AMP via their SBDβs. This structural alignment serves to highlight the inaccessibility of BiP:ADP(Thr518) for FICD engagement and AMPylation.

**Supplementary Movie 3: DeAMPylation competency is modulated by Glu234 flexibility**. Polder OMIT maps covering catalytically important residues and regions within the active site of both the state 1 (deAMPylation competent) and state 2 (deAMPylation incompetent) complexes are shown. In later frames, superposition of the two structures highlights the lack of a stably positioned catalytic water molecule directly in-line for nucleophilic attack into the Pα-Oγ(Thr518) phosphodiester bond in the state 2 complex. This is likely a result of the altered Glu234 conformation in the state 2 complex causing a shift in the position of the $Mg^{2+}$ coordination complex. Hydrogen bonds are annotated as pink dashed lines.
